# DNA methylation in *Ensifer* species during free-living growth and during nitrogen-fixing symbiosis with *Medicago* spp.

**DOI:** 10.1101/2021.03.08.434416

**Authors:** George C. diCenzo, Lisa Cangioli, Quentin Nicoud, Janis H.T. Cheng, Matthew J. Blow, Nicole Shapiro, Tanja Woyke, Emanuele G Biondi, Benoît Alunni, Alessio Mengoni, Peter Mergaert

## Abstract

Methylation of specific DNA sequences is ubiquitous in bacteria and has known roles in immunity and regulation of cellular processes, such as the cell cycle. Using single-molecule real-time sequencing, six genome-wide methylated motifs were identified across four *Ensifer* strains, five of which were strain-specific. Only the GANTC motif, recognized by the cell cycle-regulated CcrM methyltransferase, was methylated in all strains. In actively dividing cells, methylation of GANTC motifs increased progressively from the *ori* to *ter* regions in each replicon, in agreement with a cell cycle-dependent regulation of CcrM. In contrast, there was near full genome-wide GANTC methylation in the early stage of symbiotic differentiation. This was followed by a moderate decrease in the overall extent methylation and a progressive decrease in chromosomal GANTC methylation from the *ori* to *ter* regions in later stages of differentiation. We interpret these observations as evidence of dysregulated and constitutive CcrM activity during terminal differentiation, and we hypothesize that it is a driving factor for endoreduplication of terminally differentiated bacteroids.

## INTRODUCTION

Methylation of genomic DNA is a pervasive phenomenon found in eukaryotes (Greenberg and Bourc’his, 2019; Tang et al., 2012; Zhang et al., 2018), archaea (Blow et al., 2016), and bacteria (Blow et al., 2016; Sánchez-Romero et al., 2015). The biological roles of DNA methylation are most extensively studied in mammals, where it contributes to normal development and disease via its impact on gene expression (Gopalakrishnan et al., 2008). In bacteria, DNA methylation is best known for its role in restriction-modification (R-M) systems that are thought to provide defence against phage infection and limit horizontal gene transfer through the degradation of invading non-methylated DNA (Vasu and Nagaraja, 2013). Several methyltransferases (MTases) of R-M systems have also been implicated in phase variation in pathogens through modulating gene expression (Atack et al., 2018). A recent study of over 200 bacterial and archaeal species identified orphan MTases not belonging to R-M systems in nearly half of the genomes (Blow et al., 2016). To date, biological functions have been attributed to very few orphan MTases, namely, the Dam MTase of the *γ*-Proteobacteria and the CcrM MTase of the *α*-Proteobacteria (Adhikari and Curtis, 2016). The Dam MTase of *Escherichia coli* is notable for its role in regulation of DNA replication (Campbell and Kleckner, 1990; Kang et al., 1999) and DNA repair (Lahue et al., 1989) by modulating the activity of other DNA-binding proteins. The CcrM MTase was first identified in *Caulobacter crescentus* (Zweiger et al., 1994), with homologs since identified in diverse *α*-Proteobacteria (Brilli et al., 2010; Wright et al., 1997). CcrM activity was shown to be cell cycle regulated in *C. crescentus* and *Agrobacterium tumefaciens* (Kahng and Shapiro, 2001; Zweiger et al., 1994), leading to methylation of its cognate DNA motif (the pentanucleotide GANTC) specifically during a short period at the end of DNA replication. This leads to a switching of GANTC sites between fully methylated (methylated on both strands) and hemi-methylated (methylated only on the template strand) as a result of DNA replication (Kozdon et al., 2013), which serves to modulate gene expression in a cell cycle-dependent fashion (Fioravanti et al., 2013; Gonzalez et al., 2014; Gonzalez and Collier, 2013). Over- and under-expression of *ccrM* results in defects in DNA replication and cell division (Gonzalez and Collier, 2013; Kahng and Shapiro, 2001; Wright et al., 1997), while its complete loss is lethal under some conditions.

The rhizobia are a polyphyletic group of *α*-Proteobacteria and *β*-Proteobacteria that can both live free in the soil and enter into an endosymbiotic interaction with legumes (Wang and Young, 2019). This interaction begins following an exchange of signals between the free-living partners (Oldroyd, 2013), and it culminates in the formation of a new organ known as a root nodule within which the cytoplasm of plant cells contain thousands of N_2_-fixing bacteria called bacteroids. Bacteroid formation results in the differential expression of more than a thousand genes (Barnett et al., 2004; Roux et al., 2014) and global changes in cellular metabolism (diCenzo et al., 2020). In legumes of the Inverted-Repeat Lacking Clade (IRLC) and the Dalbergioid clade of the family *Papilionoideae*, bacteroid development involves an additional process of terminal differentiation (Czernic et al., 2015; Mergaert et al., 2006); in other legume clades, bacteroid differentiation is less pronounced and is reversible. Terminal bacteroid development, in contrast to reversible bacteroid formation, involves cell enlargement (bacteroids are 5- to 10-fold longer than their free-living counterparts) and genome endoreduplication (resulting in up to 24 copies of the genome per cell), indicative of a cell cycle transition occurring during differentiation (Mergaert et al., 2006). Indeed, the correct expression of cell cycle regulators in *Ensifer* (syn. *Sinorhizobium*) *meliloti*, a symbiont of *Medicago* species of the IRLC, is essential for the formation of functional bacteroids (Kobayashi et al., 2009; Pini et al., 2013), while over-expression of CcrM or disruption of the master cell cycle regulator CtrA can give rise to bacteroid-like morphology in free-living cells (Pini et al., 2015; Wright et al., 1997). Additionally, mutants in the *E. meliloti* cell cycle regulators *divJ*, *cbrA*, and *cpdR1*, encoding three negative regulators of CtrA, form non-functional nodules in which bacteroids do not differentiate properly (Gibson et al., 2006; Kobayashi et al., 2009, p. 1; Pini et al., 2013), and genes encoding several cell cycle regulators (including CcrM) are strongly downregulated in bacteroids (Roux et al., 2014). The differentiation and cell cycle switch of bacteroids is controlled by the legume host through the production of a large family of peptides, known as Nodule-specific Cysteine-Rich (NCR) peptides (Farkas et al., 2014; Penterman et al., 2014; Van de Velde et al., 2010).

Multiple studies have provided evidence that changes in the methylation status of the DNA of legume nodule cells contributes to symbiotic development (Nagymihaly et al., 2017; Pecrix et al., 2018; Satgé et al., 2016). Conversely, it remains unknown if methylation of rhizobium DNA contributes to the regulation of N_2_-fixation or bacteroid development. We are aware of only one study (Davis-Richardson et al., 2016) comparing DNA methylation of a rhizobium (*Bradyrhizobium diazoefficiens* USDA110) between free-living and symbiotic states (soybean nodules), and comparing these changes with differential expression data. Intriguingly, the authors identified a DNA motif that was methylated specifically in bacteroids (Davis-Richardson et al., 2016). However, no clear evidence was presented that methylation of this (or any other) motif is involved in transcriptional regulation, and the number of genes both differentially expressed and differentially methylated in bacteroids did not appear to be different than expected by chance. While these data may suggest that DNA methylation does not play a major role in regulating N_2_-fixation by rhizobia, they do not address the role of DNA methylation in terminal bacteroid differentiation, as *B. diazoefficiens* undergoes reversible differentiation in soybean nodules (Barrière et al., 2017).

Here, we use Pacific Biosciences Single-Molecule Real-Time (SMRT) sequencing to detect genome-wide patterns of DNA methylation in four strains belonging to the genus *Ensifer*. Our results indicate that DNA methylation is poorly conserved across the genus and suggests that DNA methylation is not a major mechanism of regulating gene expression in these organisms aside from cell cycle control. However, analysis of bacteroid samples led us to hypothesize that constitutive activation of the CcrM MTase may be a contributing factor driving terminal differentiation.

## RESULTS

### The methylomes of the genus *Ensifer*

Our experimental design, as summarized in the Materials and Methods and **Figure S1**, was developed to support an investigation into multiple potential roles of DNA methylation in plant-associated bacteria from the genus *Ensifer* through the use of SMRT sequencing. This was accomplished by: i) including DNA samples isolated from phylogenetically diverse wild-type strains, ii) examining DNA methylation in a single strain across multiple conditions (exponential phase growth versus stationary phase; growth with sucrose versus growth with succinate), iii) investigating the impact of a large-scale genome reduction on DNA methylation patterns, and iv) isolating DNA from bacteroids purified from legume nodules.

We began with base modification analyses of four wild-type strains from three species, including three nodule-forming strains (*E. meliloti* Rm2011, *E. meliloti* FSM-MA, *E. fredii* NGR234) and one plant-associated, non-symbiotic strain (*E. adhaerens* OV14). To ensure consistency, all strains were grown to mid-exponential phase in a common minimal medium with succinate as the carbon source. A total of six methylated motifs were identified, of which five were m6A modifications and one was a m4C modification (**Table 1**). Five of the six motifs were methylated specifically in one strain. Only the GANTC motif, recognized by the highly conserved cell cycle-regulated CcrM methyltransferase (Wright et al., 1997; Zweiger et al., 1994), was methylated in all four strains. To further examine the conservation, or lack thereof, of DNA modification across the genus *Ensifer*, we examined the distribution of methyltransferases in the model species *E. meliloti*. Based on gene annotations, we identified 24 genes encoding putative MTases in a previous pangenome analysis of 20 *E. meliloti* strains (**Table S1**) (diCenzo et al., 2019). Of these 24 genes, only one (*ccrM*) was found in all 20 strains, while four were found in two strains and 19 were found in a single strain. These results suggest that DNA methylation is unlikely to play a biologically significant role in the genus *Ensifer* aside from cell cycle control via CcrM-mediated methylation, and phage defence.

**Table 1.**
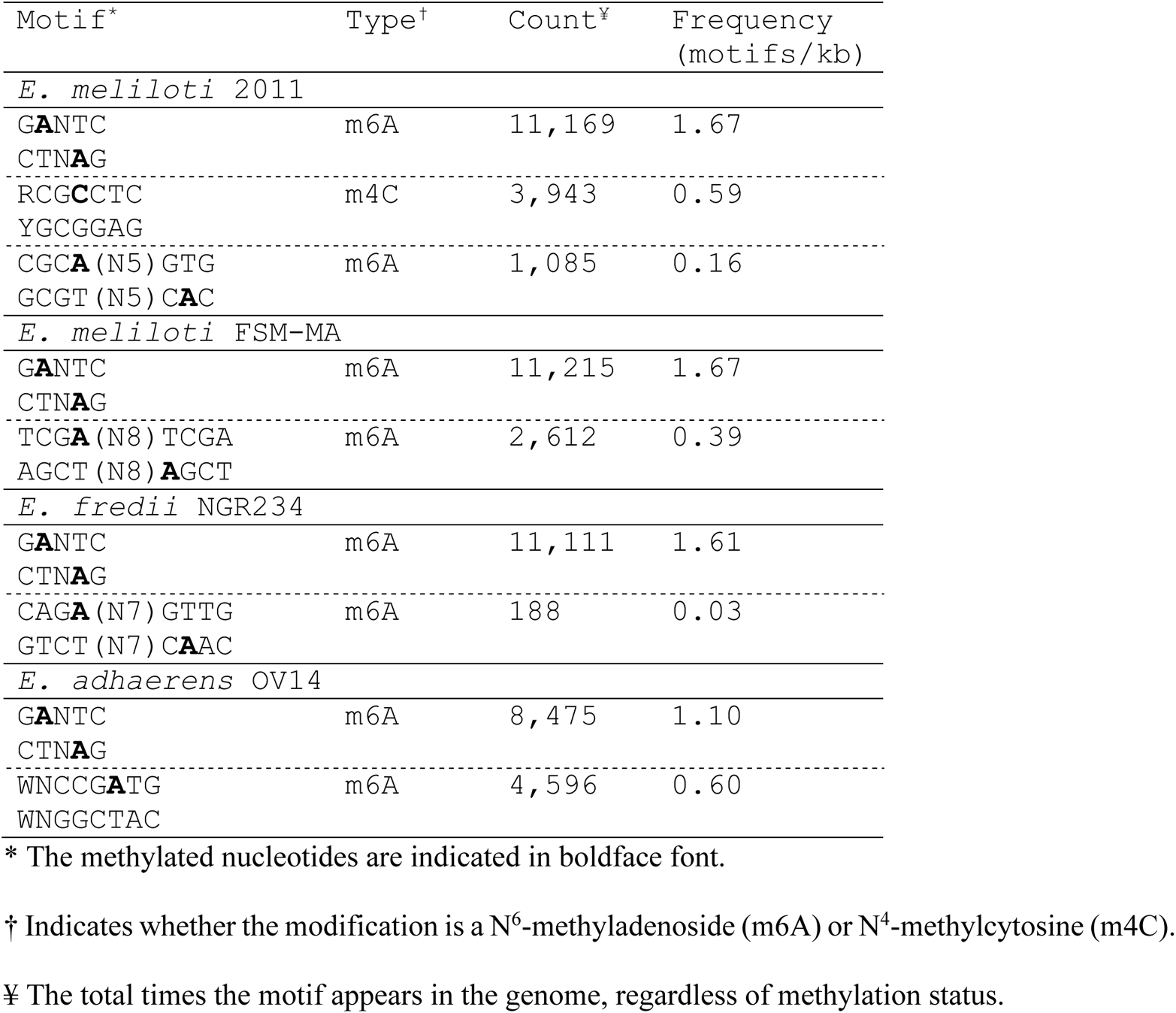
Methylated motifs identified in this study.

In support of most DNA methylation not having a regulatory function in the genus *Ensifer*, none of the motifs methylated in *E. meliloti* Rm2011 were enriched in the promoter regions of genes previously shown to be differentially expressed when grown with glucose vs. succinate (diCenzo et al., 2017). Similarly, except for the GANTC motif as discussed below, no global effect of carbon source (sucrose [glycolytic] vs. succinate [gluconeogenic]) was observed on the DNA methylation pattern of *E. meliloti* Rm2011 (**Figure S2**). Similarly no global differences in DNA methylation were detected between *E. meliloti* Rm2011 and RmP3496, a Rm2011 derivative lacking the pSymA and pSymB replicons that together account for 45% of the genome content of *E. meliloti* (diCenzo et al., 2014) (**Figure S3**).

### Cell cycle regulation by the CcrM methyltransferase

A progressive increase in the extent of methylation (herein defined as the estimated fraction of reads mapping to a motif that were methylated) of GANTC sites was observed from the *ori* to *ter* regions of the chromosomes of all four strains during mid-exponential growth (**Figures 1, S4-S6**). There was a local drop in GANTC methylation around the 1.5 Mb mark in the *E. meliloti* FSM-MA chromosome (**Figure S4**); however, this was seen in only two of three replicates and corresponded to a region of high sequencing depth (**Figure S7**), suggesting the result is a sequencing artefact. In contrast to exponential phase cultures, GANTC sites displayed near full methylation (averaging ~ 95%) across the genome during early stationary phase, while all other motifs displayed near full methylation (averaging 95-99%) across the genome regardless of growth state (**Figures 1, S4-S6**). The observed pattern of GANTC methylation indicates a progressive switch from full to hemi-methylated states as DNA replication proceeds (model provided as **Figure S8**), confirming that the CcrM methyltransferase of the family *Rhizobiaceae* is cell cycle regulated as demonstrated in *C. crescentus* (Kozdon et al., 2013; Mohapatra et al., 2014; Zweiger et al., 1994). Interestingly, the genome-wide pattern of GANTC methylation displayed a smaller variation in the extent of methylation from the *ori* to *ter* regions in *E. meliloti* Rm2011 when grown with sucrose compared to succinate as the carbon source (**Figure S2**). While this observation could suggest metabolic regulation of CcrM activity, we instead hypothesize, as displayed in **Figure S8A**, that it is due to DNA replication being initiated later in the cell cycle when *E. meliloti* is provided sucrose, as recent observations showed that central carbon metabolism influences the rate of DNA polymerase processivity and timing of DNA replication initiation in *Bacillus subtilis* (Nouri et al., 2018).

**Figure 1.**
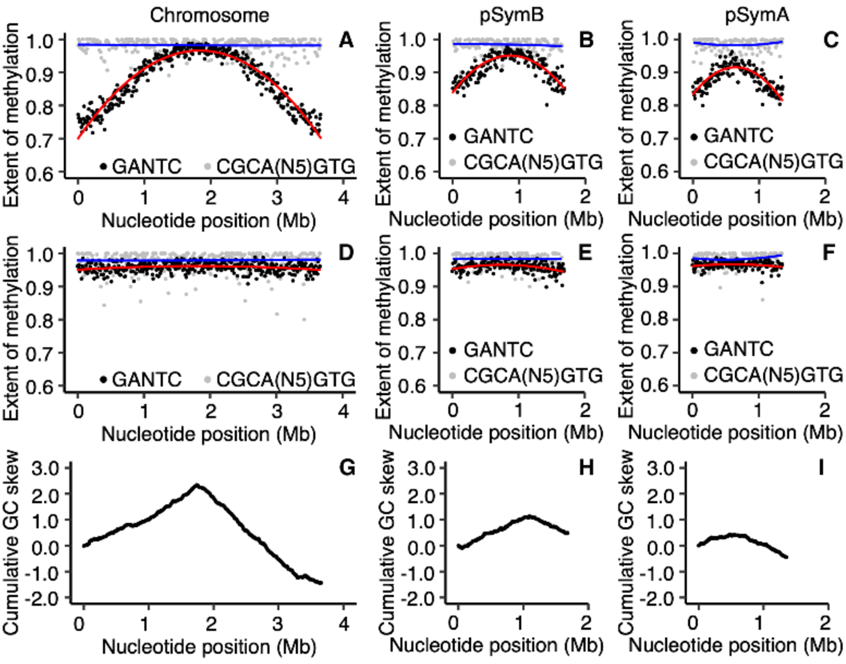
Genome-wide DNA methylation of *E. meliloti* Rm2011. (**A**-**F**) The extent of methylation is shown, using a 10 kb sliding window, of GANTC sites (black) and CGCA(N_5_)GTG sites (grey) across the chromosome (**A**,**D**), pSymB (**B**,**E**), and pSymA (**C**,**F**) replicons of exponential phase (**A**-**C**) or early stationary phase (**D**-**F**) *E. meliloti* Rm2011. Averages from three biological replicates are shown. The red (GANTC) and blue (CGCA(N_5_)GTG) lines are polynomial regression lines calculated in R using the “rlm” method and the formula “y~poly(x,2)”. (**G**-**I**) Cumulative GC skews, shown using a 10 kb sliding window, across the *E. meliloti* Rm2011 chromosome (**G**), pSymB (**H**), and pSymA (**I**) replicons.

Notably, the GANTC methylation pattern differed across replicons within each genome. For example, in *E. meliloti* Rm2011 the extent of GANTC methylation ranged from 0.80 to 0.98 for pSymB and 0.80 to 0.96 for pSymA, while for the chromosome the range was from 0.71 to 0.99 (**Figure 1**). This result suggests that replication of each replicon is asynchronous, with replication of the secondary replicons being initiated later in the cell cycle than that of the chromosome (model provided as **Figure S8**).

A previous study identified 462 cell cycle-regulated genes in *E. meliloti* through RNA-sequencing of synchronized cell populations (De Nisco et al., 2014), which were classified into six groups based on the timing of their expression. We identified 111 cell cycle-regulated genes, belonging to 78 transcripts, that contained at least one GANTC in the predicted promoter regions (defined as the 125 bp upstream of the transcript; **Dataset S1**), and distribution of these 111 genes across the six cell cycle gene expression groups was unbiased (De Nisco et al., 2014). As these 111 genes are both cell cycle regulated and contain a GANTC site, they represent an initial candidate CcrM regulon in *E. meliloti* although further work is required to validate the CcrM regulon.

We found it striking that the *E. adhaerens* OV14 genomes had 2,636 to 2,740 fewer GANTC sites than the other three strains, despite having the largest genome size. Normalized by genome length, there are 1.10 GANTC sites per kb in the *E. adhaerens* OV14 genome (**Table 1**), which is similar to the 1.12 GANTC sites per kb in *C. crescentus* NA1000. In contrast, the three legume symbionts contained more than 1.60 GANTC sites per kb across their genomes (**Table 1**). This result prompted us to examine the frequency and distribution of GANTC sites across 157 *Ensifer* genomes. As defined previously (Fagorzi et al., 2020), the genus *Ensifer* can be broadly sub-divided into two monophyletic clades; the “symbiotic” clade (113 strains) in which nearly all strains are legume symbionts, and the “non-symbiotic” clade (44 strains) in which nearly all strains are non-symbionts (**Figure 2A**). Consistent with previous results (Gonzalez et al., 2014), GANTC sites occurred less frequently in all genomes (0.90 to 1.83 GANTC sites per kb) than expected in a random sequence of nucleotides (~ 3.5 GANTC sites per kb). Moreover, GANTC sites were ~ 2-fold more common in intergenic regions than in coding regions (**Figure 2B**). Strikingly, there was a strong and statistically significant difference (*p*-value < 1 × 10^−10^; two-sample *t*-test) in the frequency of GANTC sites across the genomes of strains belonging to the symbiotic clade compared to strains of the non-symbiotic clades (**Figure 2B**), with an overall average of 1.70 and 1.06 GANTC sites per kb in the symbiotic and non-symbiotic clades, respectively. The difference in the frequency of GANTC sites between the two clades could not be explained by differences in the GC content of these organisms, as both clades had an average GC content of 61.9% (**Figure S9**), suggesting that the difference reflects underlying differences in the evolution, and possibly the biology, of these two clades.

**Figure 2.**
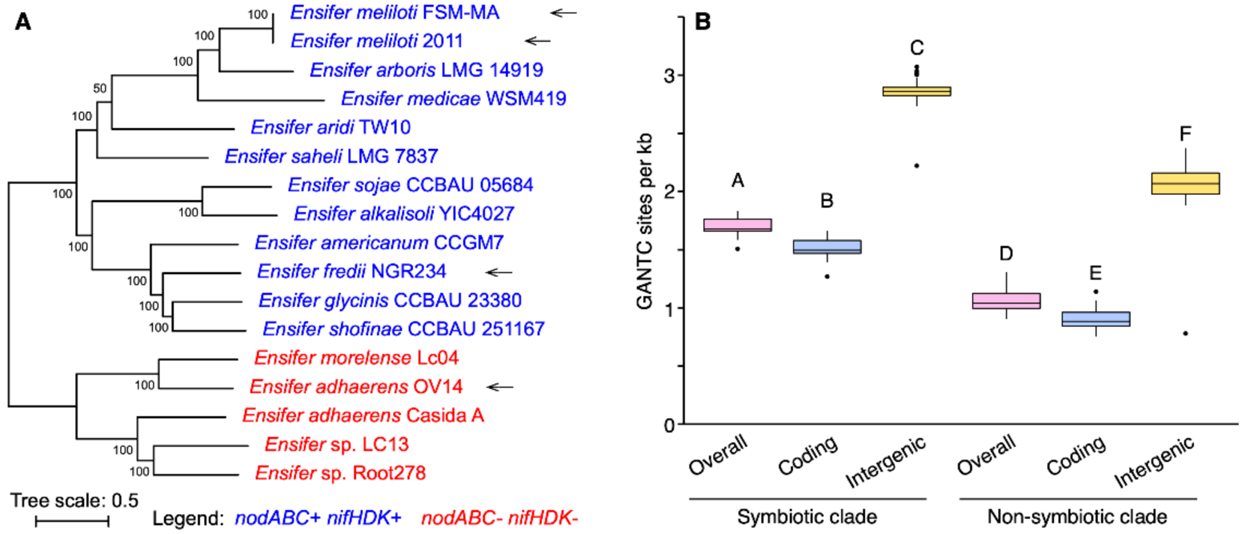
GANTC frequency in the genus *Ensifer*. (**A**) An unrooted maximum likelihood phylogeny of 17 representative *Ensifer* strains. The phylogeny represents the bootstrap best tree following 100 bootstrap replicates, prepared on the basis of the concatenated nucleotide alignments of 1566 core genes. Values represent the bootstrap support. N_2_-fixing legume symbionts were identified by the presence of the symbiotic genes *nodABC* and *nifHDK*. They are indicated in blue, while red denotes non-symbiotic strains. The four wild-type strains used in this study are indicated with arrows. (**B**) Box plots summarizing the frequency of GANTC sites (presented as GANTC sites per kb) in 157 *Ensifer* strains is shown. The monophyletic “symbiotic” and “non-symbiotic” clades as defined previously (Fagorzi et al., 2020), are represented by 113 and 44 genomes respectively. The densities of GANTC sites across the entire genome (pink), within coding regions (blue), and within intergenic regions (yellow) are shown. Statistically different values (*p* < 0.05) are denoted by uppercase letters as determined by a one-way ANOVA followed by a Tukey’s HSD post hoc test.

### Contributions of DNA methylation to bacteroid differentiation

The only previously published study to examine the role of rhizobium DNA methylation during symbiosis using SMRT sequencing did so in a symbiosis where the bacteria do not undergo terminal differentiation (Barrière et al., 2017; Davis-Richardson et al., 2016). To evaluate whether DNA methylation potentially contributes to regulation of terminal differentiation, we determined the DNA methylation patterns of *E. meliloti* Rm2011 and *E. meliloti* FSM-MA bacteroids purified from whole *Medicago sativa* nodules. *E. meliloti* FSM-MA bacteroids were additionally purified from *Medicago truncatula* nodules to determine whether the host plant influences bacteroid DNA methylation. *E. meliloti* Rm2011 bacteroids were not isolated from *M. truncatula* nodules as, unlike FSM-MA, Rm2011 forms a poor symbiosis with *M. truncatula* and produces moderately differentiated bacteroids in this host (Kazmierczak et al., 2017; Moreau et al., 2008).

Moreover, we exploited the spatially distinct developmental zones that are present in indeterminate nodules (Vasse et al., 1990), like those formed by *M. sativa* and *M. truncatula*. At the tip of these nodules, a bacteria-free meristem is present, responsible for the continuous growth of the nodule. Immediately below is the infection and differentiation zone II where nodule cells become infected and bacteria differentiate into the large, polyploid bacteroids. The tip and zone II of nodules appears white. Adjacent to the white zone II is the easily recognizable pinkish zone III (due to the presence of the oxygen-carrying leghemoglobin) where mature bacteroids fix nitrogen. This nodule tissue organization provided an opportunity to examine how DNA methylation patterns may differ between differentiating and differentiated bacteroids. To this end, *E. meliloti* Rm2011 and FSM-MA bacteroids were isolated from nodules hand-sectioned at the white-pink border; bacteroids isolated from the white sections represent the infecting and differentiating bacteroids (zone II) while those isolated from the pink sections represent the mature, hence terminally differentiated, N_2_-fixing bacteroids (zone III).

Fluorescence microscopy and flow cytometry confirmed that nodule sectioning resulted in the isolation of distinct bacteroid populations (**Figures S10-S13**). Nearly all of the bacteroids isolated from whole-nodule samples and zone III samples were enlarged and polyploid, and most were positive for propidium iodide (PI) staining as expected for terminally differentiated bacteroids (Mergaert et al., 2006). In contrast, bacteroids of the zone II samples contained a mix of cell types differing in their size, ploidy level, and PI staining. These data confirmed that the whole-nodule samples and zone III samples consisted predominately of mature N_2_-fixing bacteroids, whereas the zone II samples contained a mix of cells at various stages of bacteroid differentiation.

With the exception of the GANTC sites (described below), no global difference was observed in the methylation patterns of bacteroids versus free-living cells (**Figures 3, 4, S14-S17**). Although a lower percentage of each motif was detected as methylated in the bacteroid samples compared to free-living samples, this was correlated with lower sequencing depth (**Table S2**) and thus unlikely to be biologically meaningful. Unlike *B. diazoefficiens* USDA110 (Davis-Richardson et al., 2016), we did not identify motifs that were methylated specifically in the *E. meliloti* bacteroids. In addition, we found little evidence for any of the methylated motifs being enriched in the promoter regions of *E. meliloti* Rm2011 genes up-regulated or down-regulated in bacteroids relative to free-living cells, as identified in published transcriptomic data for *E. meliloti* Rm1021 (Barnett et al., 2004), a near-isogenic relative of strain Rm2011 also derived from the nodule isolate SU47. These data suggest that most DNA methylation is unlikely to be a significant factor in directly regulating gene expression in *E. meliloti* bacteroids.

**Figure 3.**
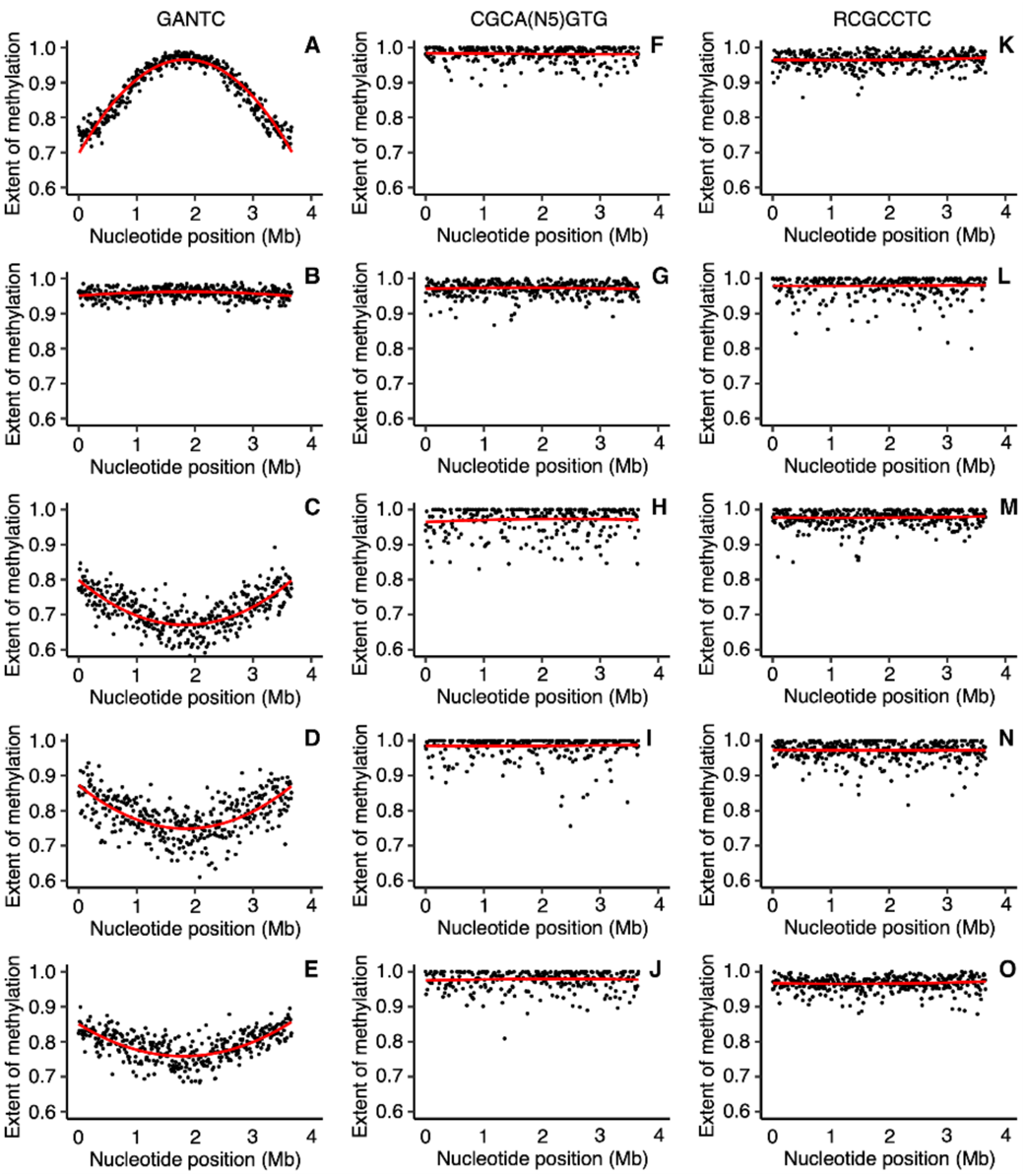
Chromosome-wide DNA methylation of *E. meliloti* Rm2011 bacteroids. The extent of methylation of (**A**-**E**) GANTC, (**F**-**J**) CGCA(N_5_)GTG, and (**K**-**O**) RCGCCTC motifs across the *E. meliloti* Rm2011 chromosome is shown using a 10 kb sliding window. Averages from three biological replicates are shown for free-living and whole nodule samples; data represents one replicate for the zone II and zone III nodule sections. (**A**,**F**,**K**) Free-living cells harvested in mid-exponential phase. (**B**,**G**,**L**) Free-living cells harvested in early stationary phase. (**C**,**H**,**M**) Bacteroids isolated from *M. sativa* zone II nodule sections. (**D**,**I**,**N**) Bacteroids isolated from *M. sativa* zone III nodule sections. (**E**,**J**,**O**) Bacteroids isolated from *M. sativa* whole nodule samples. The red lines are polynomial regression lines calculated in R using the “rlm” method and the formula “y~poly(x,2)”. Data for pSymB and pSymA are shown in Figures S14 and S15.

**Figure 4.**
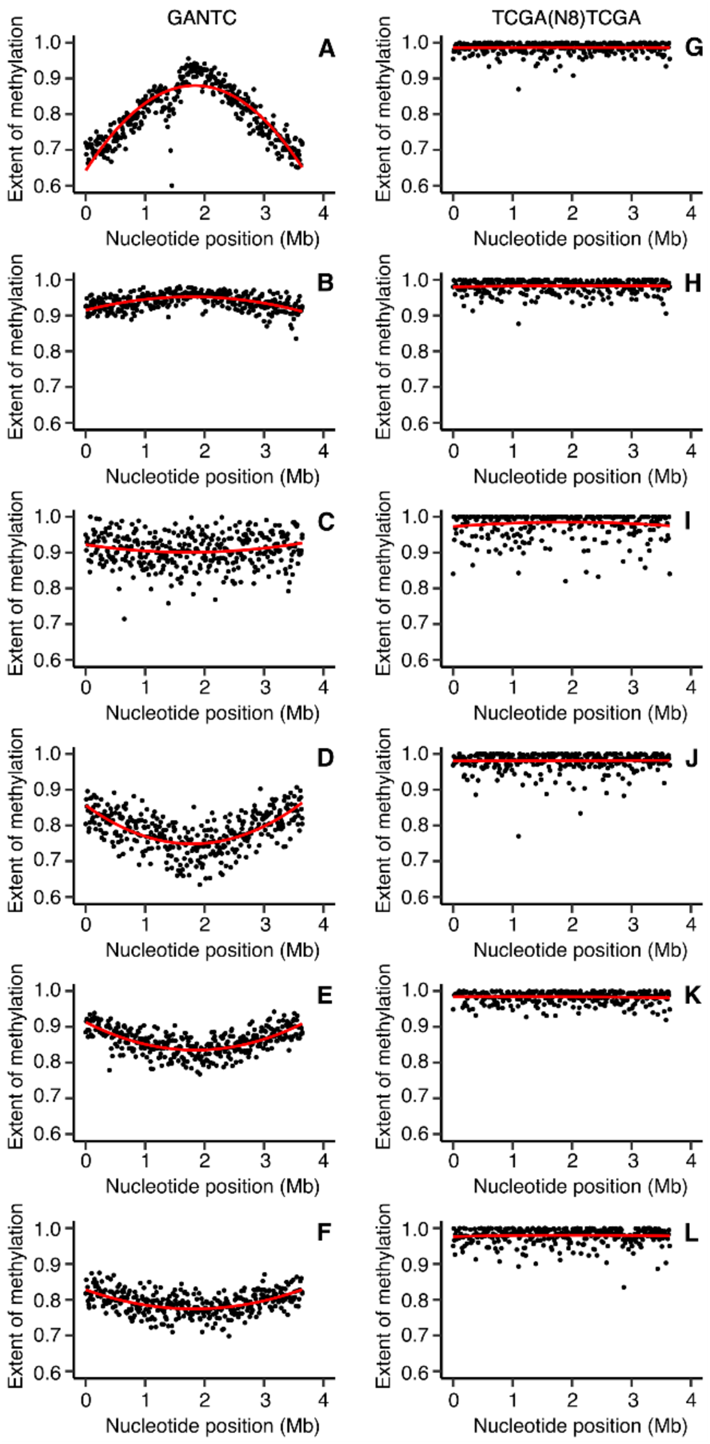
Chromosome-wide DNA methylation of *E. meliloti* FSM-MA bacteroids. The extent of methylation of (**A**-**F**) GANTC and (**G**-**L**) TCGA(N8)TCGA motifs across the *E. meliloti* FSM-MA chromosome is shown using a 10 kb sliding window. Averages from three biological replicates are shown for free-living and whole nodule samples; data represents one replicate for the zone II and zone III nodule sections. (**A**,**G**) Free-living cells harvested in mid-exponential phase. (**B**,**H**) Free-living cells harvested in early stationary phase. (**C**,**I**) Bacteroids isolated from *M. sativa* zone II nodule sections. (**D,J**) Bacteroids isolated from *M. sativa* zone III nodule sections. (**E**,**K**) Bacteroids isolated from *M. sativa* whole nodule samples. (**F**,**L**) Bacteroids isolated from *M. truncatula* whole nodule samples. The red lines are polynomial regression lines calculated in R using the “rlm” method and the formula “y~poly(x,2)”. Data for pSymB and pSymA are shown in Figures S16 and S17.

### CcrM methyltransferase activity is dysregulated during terminal differentiation

Bacteroid development involves cell enlargement and genome endoreduplication, indicative of a cell cycle transition occurring during differentiation (Mergaert et al., 2006). Indeed, expression of *ccrM* and other genes encoding cell cycle regulators vary across stages of bacteroid development and are strongly downregulated in mature nitrogen-fixing *E. meliloti* bacteroids (Roux et al., 2014). We were therefore interested in examining whether GANTC methylation by the CcrM MTase was disrupted in bacteroids. Our data revealed a surprising genome-wide pattern of GANTC methylation in *E. meliloti* Rm2011 and FSM-MA bacteroids, which differed from free-living cells in either the exponential or stationary phases of growth (**Figures 3, 4, S14-S17**). The majority of GANTC sites had moderate to high levels of methylation in zone II, zone III, or whole-nodule samples, averaging 0.71 to 0.95 across each replicon (**Figures 3, 4, S14-S17, and Tables S3, S4**). Most distinctive, a progressive decrease in the extent of chromosomal methylation of the GANTC sites was observed from the *ori* to *ter* in nearly all bacteroid samples, revealing a characteristic smiling pattern, which differs from the patterns seen in exponential (frowning pattern, i.e., a progressive increase from *ori* to *ter*) and stationary (consistent methylation) phase cells. The exception was the *E. meliloti* FSM-MA zone II bacteroid sample, which displayed a consistently high level of GANTC methylation across the genome (**Figure 4C**). This pattern, which is different from those in exponential phase cells as well as mature bacteroids, could correspond to the methylation status of an early stage of bacteroid differentiation. We did not observe the same pattern in the *E. meliloti* Rm2011 zone II samples. As noted earlier (**Figures S10-S13**), the zone II samples contain cells at various stages of differentiation. Given that terminal differentiation is associated with an up to 24-fold increase in DNA content, small increases in the proportion of cells at late stages of differentiation could mask the DNA methylation pattern of the cells at early stages of differentiation. Thus, we hypothesize that the Rm2011 zone II sample captures a later stage of differentiation than that captured by the FSM-MA zone II sample. Supporting this hypothesis, the distribution of DNA content per cell in the flow cytometry data was flatter for *E. meliloti* Rm2011 zone II bacteroids compared to *E. meliloti* FSM-MA zone II bacteroids (**Figure S18**), which suggests that the former sample represents a broader range of differentiation stages than the latter sample. In contrast to GANTC, the extent of methylation of the second m6A modified motif in each genome was consistently high, irrespective of condition or replicon (**Figures 3, 4, S14-S17, and Tables S3, S4**). Similarly, sequencing depth was consistent across the length of each replicon (**Figure S7**). These observations indicate that the changes in GANTC methylation patterns are biologically meaningful and not simply a sequencing artefact.

To further explore changes in CcrM activity during bacteroid differentiation, we took advantage of a collection of *M. truncatula* mutant plant lines (*dnf1*, *dnf2*, *dnf4*, *dnf5*, *dnf7*) whose nodules contain bacteria blocked at various stages of differentiation (Bourcy et al., 2013; Domonkos et al., 2013; Horváth et al., 2015; Kim et al., 2015; Lang and Long, 2015; Starker et al., 2006; Wang et al., 2010). Microscopy and flow cytometry data was consistent with past work (Lang and Long, 2015) showing that bacteroids were blocked at the earliest to latest stages of differentiation in mutant plant lines in the order *dnf1* à *dnf5* à *dnf2* à *dnf7* à *dnf4* (**Figure 5**). Nodule bacteria of *M. truncatula dnf1* mutant plants were small with one or two haploid genome copies per cell (i.e., ploidy level = 1 or 2) (**Figures 5A, 5G**), suggesting that the cell population was dominated by actively dividing cells that had not yet begun differentiation. Indeed, the GANTC methylation pattern of these cells (**Figures 5M, S19, S20**) resembled the frowning pattern of exponentially growing free-living cells (**Figure 4A**). Although the nodule bacteria of *M. truncatula dnf5* mutant plants were also small and undifferentiated into bacteroids, the majority of cells had a ploidy level of one (**Figures 5B, 5H**), suggesting these cells had ceased replication but had not yet begun the process of endoreduplication. GANTC methylation was consistently high across the chromosome of bacteria purified from *dnf5* nodules (**Figure 5N**), similar to stationary phase free-living cells (**Figure 4B**) and indicating that terminal differentiation is preceded by full GANTC methylation.

**Figure 5.**
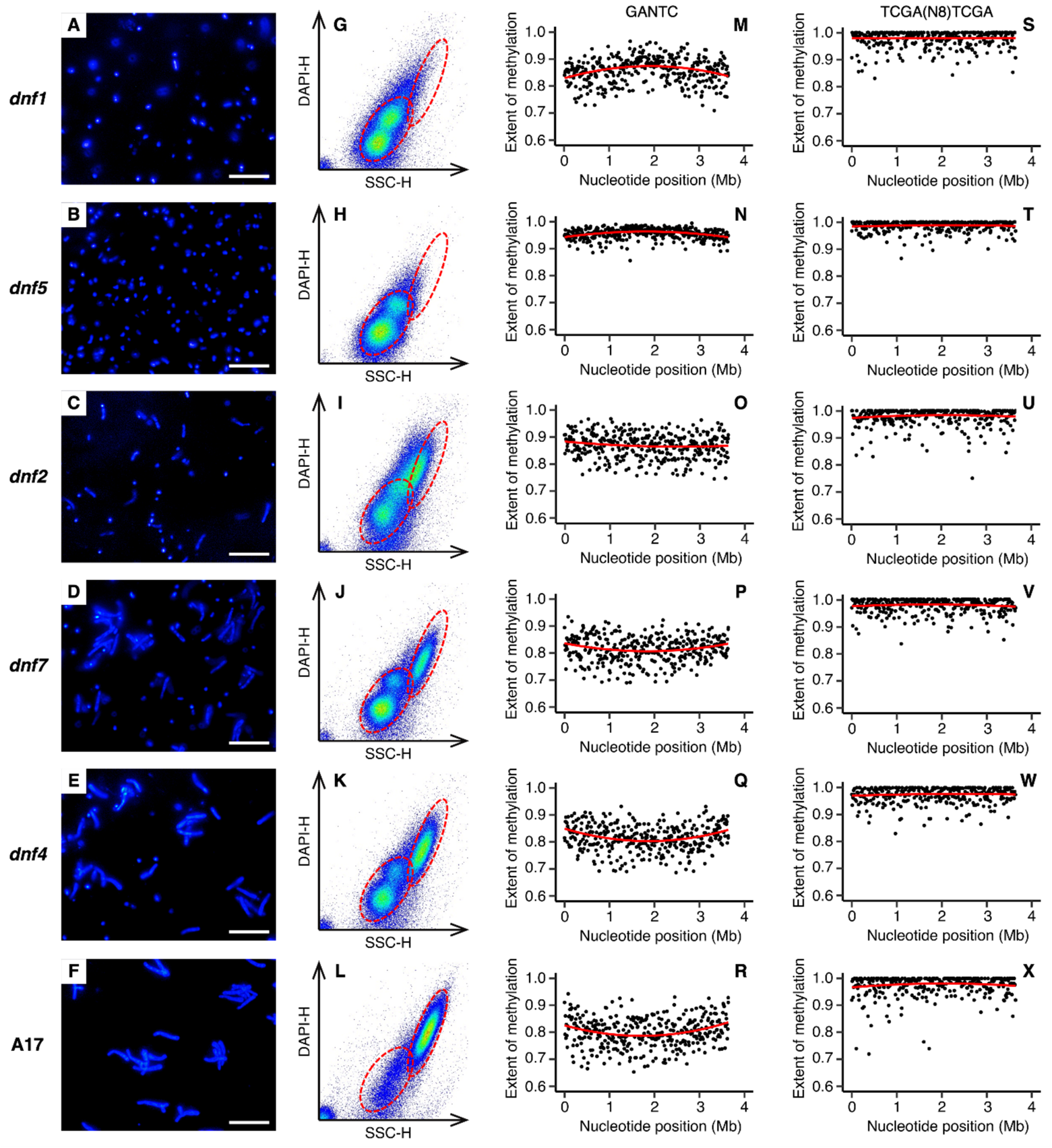
Bacteroid morphology and chromosomal GANTC methylation in *E. meliloti* bacteroids purified from *M. truncatula dnf* mutant nodules. Data is shown for *E. meliloti* FSM-MA bacteroids purified from (**A**,**G**,**M**,**S**) *M. truncatula dnf1* mutant nodules, (**B**,**H**,**N**,**T**) *M. truncatula dnf5* mutant nodules, (**C**,**I**,**O**,**U**) *M. truncatula dnf2* mutant nodules, (**D**,**J**,**P**,**V**) *M. truncatula dnf7* mutant nodules, (**E**,**K**,**Q**,**W**) *M. truncatula dnf4* mutant nodules, and (**F**,**L**,**R**,**X**) *M. truncatula* A17 wild-type nodules. (**A**-**F**) Micrographs of *E. meliloti* FSM-MA bacteroids stained with the DNA binding dye DAPI. The scale bar represents 30 µm. (**G**-**L**) Pseudo-coloured scatterplots displaying the cell morphology (X-axis) and DNA content (Y-axis) of *E. meliloti* FSM-MA bacteroids, as determined based on flow cytometry analysis of DAPI stained cells. The red dashed ellipses indicate the position of undifferentiated bacteria as in culture (not shown) or in the *dnf1* mutant nodules (lower left ellipse) or fully mature bacteroids as in the A17 wild-type nodules (top right ellipse). (**M**-**X**) The extent of methylation of (**M**-**R**) GANTC or (**S**-**X**) TCGA(N_8_)TCGA motifs across the *E. meliloti* FSM-MA chromosome, shown using a 10 kb sliding window. The red lines are polynomial regression lines calculated in R using the “rlm” method and the formula “y~poly(x,2)”. Data for pSymB and pSymA are shown in Figures S19 and S20.

The nodule bacteria purified from *M. truncatula dnf2* mutant nodules were a mix of undifferentiated and partially differentiated bacteroids, which were polyploid to an extent similar to bacteroids purified from wild-type A17 nodules (**Figure 5I** compared to **Figure 5L**); however, their cell size was much smaller (**Figure 5C** compared to **Figure 5F**). This was similar to differentiating bacteroids purified from *M. truncatula* and *M. sativa* zone II nodule sections, many of which had high ploidy without a corresponding increase in cell size (**Figures S11, S13**). The GANTC methylation pattern of bacteroids from *dnf2* nodules (**Figure 5O**) was also similar to that of zone II nodule sections. There was a consistently high extent of GANTC methylation across the chromosome averaging 0.870, which was less than that of bacteroids purified from *dnf5* nodules (0.956) but higher than that of bacteroids purified from wild-type A17 nodules (0.804) (**Table S5**), and without the smiling pattern. The nodule bacteria purified from *M. truncatula dnf7* and *dnf4* nodules also contained a mix of undifferentiated bacteria and fully differentiated bacteroids (**Figures 5D, 5E, 5J, 5K**), with the number of undifferentiated bacteria greater in *dnf7* nodules compared to *dnf4* nodules. The GANTC methylation pattern of bacteroids purified from *dnf7* and *dnf4* nodules was similar to that of bacteroids purified from A17 nodules (**Figures 5P-Q**). Overall, we interpret the data from bacteroids purified from section nodules and *M. truncatula dnf* mutant nodules as suggesting that CcrM is dysregulated during terminal bacteroid differentiation and that CcrM is constitutively active during endoreduplication.

### Chromosome, pSymB, and pSymA sequencing depth are unequal in *E. meliloti* bacteroids

We noticed that in each bacteroid sample, the average extent of GANTC methylation for the chromosomes of the two strains were lower (by 0.04 to 0.13) than that of pSymA or pSymB, and unlike the chromosome, the extent of GANTC methylation was relatively constant across pSymA and pSymB (**Figures S14-S17** compared to **Figures 3 and 4**). These results suggest that, unlike in free-living cells, replication of the three replicons is not well coordinated during terminal differentiation. In agreement with this hypothesis, the mean sequencing depth across pSymA and pSymB was on average ~ 33% lower than that of the chromosome in all replicates of the *E. meliloti* whole-nodule bacteroid samples (**Table 2**). Similarly, the mean sequencing depth across pSymA and pSymB was on average ~ 23% lower than that of the chromosome for the polyploid bacterial cell populations purified from *M. truncatula dnf2*, *dnf7*, and *dnf4* mutant nodules, but not for the haploid/diploid bacterial cell populations purified from *M. truncatula dnf1* and *dnf5* mutant nodules (**Table 2**). Assuming sequencing depth is correlated with copy number, this observation suggests that *E. meliloti* bacteroids carry approximately two copies each of pSymA and pSymB per three copies of the chromosome.

**Table 2.**
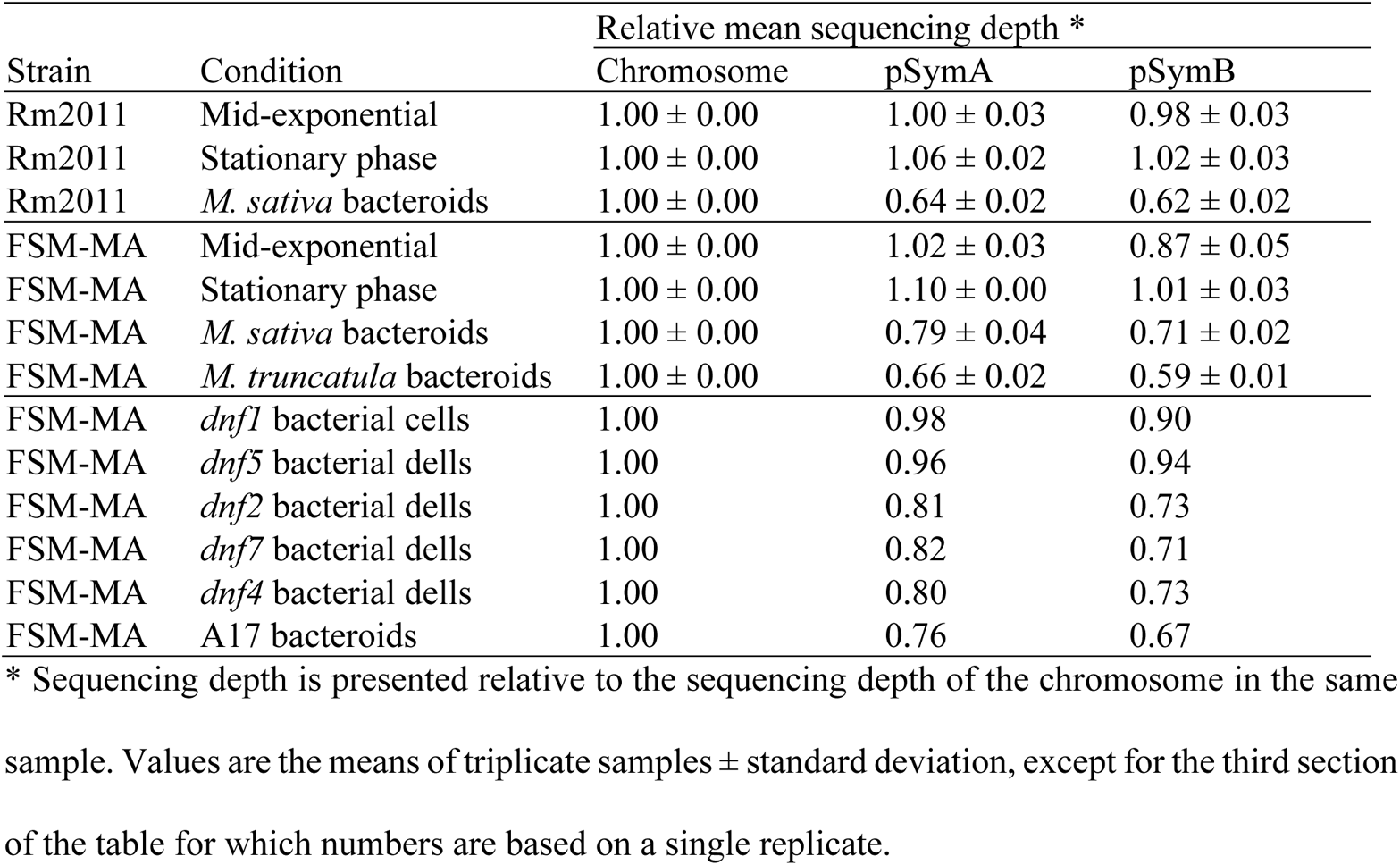
Relative sequencing depth of each *E. meliloti* replicon.

| Strain | Condition | Relative mean sequencing depth * |  |  |
| --- | --- | --- | --- | --- |
|  |  | Chromosome | pSymA | pSymB |
| Rm2011 | Mid-exponential | 1.00 ± 0.00 | 1.00 ± 0.03 | 0.98 ± 0.03 |
| Rm2011 | Stationary phase | 1.00 ± 0.00 | 1.06 ± 0.02 | 1.02 ± 0.03 |
| Rm2011 | *M. sativa* bacteroids | 1.00 ± 0.00 | 0.64 ± 0.02 | 0.62 ± 0.02 |
| FSM-MA | Mid-exponential | 1.00 ± 0.00 | 1.02 ± 0.03 | 0.87 ± 0.05 |
| FSM-MA | Stationary phase | 1.00 ± 0.00 | 1.10 ± 0.00 | 1.01 ± 0.03 |
| FSM-MA | *M. sativa* bacteroids | 1.00 ± 0.00 | 0.79 ± 0.04 | 0.71 ± 0.02 |
| FSM-MA | *M. truncatula* bacteroids | 1.00 ± 0.00 | 0.66 ± 0.02 | 0.59 ± 0.01 |
| FSM-MA | *dnf1* bacterial cells | 1.00 | 0.98 | 0.90 |
| FSM-MA | *dnf5* bacterial dells | 1.00 | 0.96 | 0.94 |
| FSM-MA | *dnf2* bacterial dells | 1.00 | 0.81 | 0.73 |
| FSM-MA | *dnf7* bacterial dells | 1.00 | 0.82 | 0.71 |
| FSM-MA | *dnf4* bacterial dells | 1.00 | 0.80 | 0.73 |
| FSM-MA | A17 bacteroids | 1.00 | 0.76 | 0.67 |

\* Sequencing depth is presented relative to the sequencing depth of the chromosome in the same sample. Values are the means of triplicate samples ± standard deviation, except for the third section of the table for which numbers are based on a single replicate.

## DISCUSSION

In this study, we examined the genome-wide DNA methylation patterns in the free-living cells of four *Ensifer* strains, and in bacteroids of two *E. meliloti* strains, and detected a total of six methylated motifs. We were able to predict cognate MTases for most of the motifs based on genome annotations, the exception being the WNCCGATG motif of *E. adhaerens* OV14. The CGCA(N_5_)GTG motif of *E. meliloti* Rm2011 is presumably methylated by Smc02296 (HsdM), a predicted m6A MTase belonging to the HsdRSM type I R-M system that is known to be functional and reduce transformation efficiency (Brumwell et al., 2019; Ferri et al., 2010). The RCGCCTC motif of *E. meliloti* Rm2011 is possibly methylated by Smc03763, a predicted cysteine-specific MTase located upstream of the gene *vsr* that putatively encodes a very short patch repair protein. Neither of these proteins are found in the other three strains examined here. The motifs TCGA(N_8_)TCGA of *E. meliloti* FSM-MA and CAGA(N_7_)GTTG of *E. fredii* NGR234 are likely methylated by SMB554_16155 and NGR_c01340, respectively, which are 88% identical at the amino acid level. Homologs of these two proteins are not found in the other two strains.

Except for GANTC, each methylated motif was detected as methylated only in a single strain. Moreover, MTases, apart from CcrM, are not conserved among *E. meliloti* strains. The lack of conservation suggests that most DNA methylation does not have a major regulatory role in the genus *Ensifer*, aside from its role in cell cycle regulation. Supporting this conclusion, no motif was enriched in the promoter regions of symbiosis, carbon source, or cell cycle-regulated genes, and we did not detect any motifs that were methylated specifically in bacteroids. However, we cannot rule out that one or more methylated motifs may influence specific gene expression during free-living growth, differentiation, or N_2_-fixation through extended motifs or proximity to other promoter elements, similar to the interplay between CcrM and GcrA during cell cycle regulation in *C. crescentus* (Fioravanti et al., 2013; Haakonsen et al., 2015).

As previously published studies have provided evidence for a cell cycle transition occurring during terminal bacteroid differentiation (Mergaert et al., 2006), we were particularly interested in CcrM, a cell cycle-regulated MTase that is broadly conserved in the *α*-Proteobacteria, and its cognate DNA motif, GANTC. By identifying GANTC sites in the promoter regions of a previously determined set of 462 cell cycle-regulated genes (De Nisco et al., 2014), we defined a candidate CcrM regulon in *E. meliloti* consisting of 111 genes. However, further studies are required to better delineate the CcrM regulon in *E. meliloti* as the presence of a GANTC site is not diagnostic of CcrM regulation; GANTC sites were found in the promoter regions of 904 transcripts that did not display cell cycle regulation, and the promoter regions of cell cycle regulated genes were not enriched in GANTC sites relative to the whole *E. meliloti* genome. Studies in *C. crescentus* suggest that the impact of the fully or hemi-methylated status of GANTC sites on gene expression is mediated, at least in part, through modulating the activity of the transcriptional regulator GcrA (Fioravanti et al., 2013; Haakonsen et al., 2015). However, not all promoter sites containing a GANTC motif are regulated by GcrA in *C. crescentus*, with the relationship dependent on an extended YGAKTCK motif and the precise position of this motif relative to other promoter elements (Haakonsen et al., 2015). Likely, CcrM-mediated gene regulation in the genus *Ensifer* is similarly dependent upon additional sequence elements beyond the GANTC motif.

Consistent with past observations (Gonzalez et al., 2014), GANTC sites were under-represented in the genomes of 157 *Ensifer* strains, particularly within coding regions. More surprising, however, was the strong difference in the frequency of GANTC sites between the previously defined (Fagorzi et al., 2020) symbiotic and non-symbiotic clades in the genus *Ensifer*, with the frequency of GANTC sites being ~ 60% higher in the symbiotic clade. As methylation of the GANTC motif by CcrM is known to influence gene expression in *C. crescentus* (Gonzalez et al., 2014), our observations suggest that CcrM has a greater impact on modulating gene expression in the symbiotic clade compared to the non-symbiotic clade. Although further work is required to understand the biological significance of the greater frequency of GANTC sites in the symbiotic clade, it is tempting to speculate it is associated with legume symbiosis.

Our data is consistent with CcrM activity differing during terminal bacteroid differentiation compared to free-living cells. The overall moderate to high rates of GANTC methylation in all *E. meliloti* bacteroid samples, coupled with the lack of a chromosome-wide pattern in the *E. meliloti* FSM-MA zone II sample, leads us to hypothesize that CcrM remains constitutively active throughout most of terminal differentiation. This hypothesis is supported by the results for nodule bacteria purified from *M. truncatula dnf* mutant nodules, which showed that differentiation is preceded by full GANTC methylation and that GANTC methylation remains high (but moderately lower) during endoreduplication followed by another moderate drop in GANTC methylation in late stages of differentiation. Considering that over-expression of CcrM can give rise to bacteroid-like morphology in free-living cells (Wright et al., 1997), we hypothesize that constitutive CcrM MTase activity is one (of potentially multiple) factor(s) driving polyploidization of bacteroids (**Figure S8**). However, further studies monitoring CcrM abundance and artificially manipulating *ccrM* expression throughout bacteroid differentiation are required to conclusively determine if CcrM is constitutively active during terminal differentiation and the importance of this activity to the promotion of endoreduplication.

CcrM activity is confined to a short window in the cell cycle since the *ccrM* gene is expressed in the late phase of genome replication (De Nisco et al., 2014) and the CcrM protein is degraded by the Lon protease prior to cell division (Wright et al., 1996). Thus, constitutive CcrM MTase activity in differentiating bacteroids could be obtained through an aberrant expression of the gene or alternatively a lack of proteolytic degradation of the CcrM protein. In agreement with the latter possibility, Lon protease was identified as a target of the NCR247 peptide (Farkas et al., 2014). It is tempting to speculate that NCR peptides like NCR247 inhibit Lon protease activity post-translationally, thereby stabilizing CcrM and triggering bacteroid differentiation. However, the CcrM MTase does not appear to remain active in fully differentiated bacteroids, with the lower GANTC methylation near the chromosomal *ter* regions suggesting that loss of CcrM MTase activity occurs slightly prior to completion of genome endoreduplication (model provided as **Figure S8**). These hypotheses are consistent with *M. truncatula* – *E. meliloti* nodule zone-specific RNA-sequencing data (Roux et al., 2014), which showed that *ccrM* expression in the root distal portion of zone II is ~ 2-fold higher than in the root proximal portion of zone II, and ~ 10-fold higher than in zone III. The ~ 10-fold difference in *ccrM* expression across nodule zones suggests to us that the level of *ccrM* expression during early stages of bacteroid differentiation is biologically significant, a prerequisite for the constitutive CcrM activity that we hypothesize.

Our analyses also provide insight into the genome replication dynamics of *E. meliloti* during free-living growth and terminal bacteroid differentiation. Notably, flow cytometry data of *E. meliloti* bacteroids purified from zone II nodule sections and *M. truncatula dnf2* nodules suggest that endoreduplication and cell enlargement largely occur subsequently, not concurrently. Genome replication might be a much faster process than cell growth or alternatively, endoreduplication might be required to drive cell enlargement. Moreover, our data are consistent with a loss of coordination of replication of the three replicons during terminal bacteroid differentiation, leading to unequal copy numbers with two copies of pSymA and pSymB per three copies of the chromosome in bacteroids. This relative change in replicon copy number occurs concomitantly with differentiation and polyploidization, as supported by the relative abundance of the replicons differing in nodule bacterial cells that have experienced endoreduplication (i.e., cells retrieved from *M. truncatula dnf2*, *dnf7*, and *dnf4* mutant nodules) but not in bacterial cells that had not yet undergone endoreduplication (i.e., cells retrieved from *M. truncatula dnf1* and *dnf5* mutant nodules). This differs from free-living cells, where copy number of the three replicons was approximately equal based on average sequencing depth. In contrast to our results, a previous comparison of the relative abundance of the three replicons in free-living *E. meliloti* Rm1021 versus bacteroids, no changes were detected using comparative genome hybridization with microarrays (Mergaert et al., 2006). We believe that the difference between our present data and the previous analysis is due to the subtlety of the differences and the lower sensitivity of the microarray hybridization method compared to high throughput sequencing.

We also observed that during free-living exponential growth, the extent of GANTC methylation at the *ori* of pSymA and pSymB is higher than at the *ori* of the chromosome, while the *ter* of the pSymA and pSymB has a slightly lower extent of GANTC methylation than the *ter* of the chromosome. As GANTC methylation occurs at a fixed stage of the cell cycle corresponding to the end of chromosome replication, our observations indicate that pSymA and pSymB replication is initiated later in the cell cycle than initiation of chromosome replication, while their replication terminates slightly before completion of chromosomal replication and the activation of CcrM (model provided as **Figure S8**). These results provide additional support for the notion of spatiotemporal regulation of DNA replication and partitioning in the multipartite *E. meliloti* genome as proposed previously (De Nisco et al., 2014; Frage et al., 2016). Similarly, replication of chromosome II of *Vibrio cholerae* is delayed relative to chromosome I, leading to the replication of these two replicons terminating at approximately the same time (Rasmussen et al., 2007). Thus, co-ordinating the timing or replication termination may be a general feature of multipartite genomes.

## MATERIALS AND METHODS

### Experimental design

The overall experimental design is summarized in **Figure S1**. Genomic DNA was isolated from four wild-type *Ensifer* strains to explore how DNA methylation varies across this genus; to allow direct comparison, the four strains were grown to mid-exponential phase in minimal medium with succinate as a carbon source. To investigate how DNA methylation patterns differ between actively dividing and non-dividing cells, genomic DNA was isolated from *E. meliloti* Rm2011 growth to either mid-exponential phase or stationary phase. Genomic DNA was isolated from *E. meliloti* Rm2011 grown to mid-exponential phase with either a glycolytic (sucrose) or gluconeogenic (succinate) carbon source to investigate whether DNA methylation may play a role in regulating carbon metabolism. Furthermore, a *E. meliloti* Rm2011 derivative lacking the pSymA and pSymB replicons (named RmP3496) was studied to gain insight into whether these replicons contribute to DNA methylation patterns in *E. meliloti* Rm2011; these strains were grown with sucrose (instead of succinate) as RmP3496 lacks the succinate transporter.

In addition to the free-living samples, *E. meliloti* bacteroid samples purified from legume nodules were collected to investigate changes in DNA methylation during bacteroid differentiation and nitrogen fixation. To do so, *E. meliloti* Rm2011 and *E. meliloti* FSM-MA bacteroids were isolated from *M. sativa* whole nodules. *E. meliloti* FSM-MA bacteroids were additionally purified from *M. truncatula* whole nodules to examine the impact of host plant on bacteroid DNA methylation patterns. *E. meliloti* Rm2011 bacteroids were only isolated from *M. sativa* nodules as, unlike FSM-MA, Rm2011 forms a poor symbiosis with *M. truncatula* (Kazmierczak et al., 2017; Moreau et al., 2008). Moreover, *E. meliloti* Rm2011 and *E. meliloti* FSM-MA bacteroids were isolated from *M. sativa* nodule sections (sectioned at the white – pink border to separate the root distal infection and differentiation zone II [white] from the root proximal nitrogen-fixing zone III [pink]) to facilitate an analysis of how DNA methylation patters differ between differentiating bacteroids and fully differentiated and nitrogen-fixing bacteroids. This was followed by isolation of *E. meliloti* FSM-MA bacteroids from whole nodules of *M. truncatula* mutant lines (*dnf1*, *dnf2*, *dnf4*, *dnf5*, *dnf7*) to investigate DNA methylation patterns in nodule bacteria blocked as various stages of differentiation.

### Bacterial strains and growth conditions

Bacterial strains used in this work are listed in **Table S6**. All strains were routinely grown on TY with 2 µM CoCl_2_ as it was required for *E. meliloti* RmP3496 (diCenzo et al., 2014). The MM9 minimal medium (diCenzo et al., 2014) consisted of the following: 40 mM MOPS, 20 mM KOH, 19.2 mM NH_4_Cl, 85.6 mM NaCl, 2 mM KH_2_PO_4_, 1 mM MgSO_4_, 0.25 mM CaCl_2_, 1 µg ml^−1^ biotin, 42 nM CoCl_2_, 38 µM FeCl_3_, 10 µM thiamine-HCl, and either 10 mM sucrose (MM9-sucrose) or 20 mM disodium succinate (MM9-succinate). Prior to inoculation of plants with *E. meliloti*, the strains were grown in YEB medium (Krall et al., 2002).

### DNA isolation from free-living cells

Overnight cultures of all strains grown in MM9-succinate or MM9-sucrose media were diluted into 10 mL of the same medium to a starting OD_600nm_ of 0.025 (0.05 for RmP3496) and incubated overnight at 30°C with shaking (130 rpm). The next day, cultures were diluted into 40 mL of the same medium in 100 mL flasks to the OD_600nm_ values as listed in **Table S7**. To obtain mid-exponential phase samples, cultures were harvested after 15.5-16 hours of growth at OD_600nm_ values between 0.37 and 0.69 (**Table S7**). To obtain stationary phase samples, cultures were harvested after 24 hours of growth at OD_600nm_ values of ~ 1.4. In all cases, cultures were streaked on TY plates to check for contamination and then centrifuged (8,200 *g*, 10 minutes, 4°C); the full 40 mL was centrifuged for mid-exponential phase cultures, whereas only 15 mL was centrifuged for stationary phase cultures. Most of the supernatant was discarded, and the pellet resuspended in the remaining volume, transferred to a 2 mL tube, centrifuged again (16,200 *g*, room temperature, one minute), and the supernatant discarded. Three biological replicates, each starting from a separate overnight culture, were performed. DNA was isolated using phenol:chloroform extractions and ammonium acetate precipitations as described elsewhere (Cowie et al., 2006), and the DNA pellets (following RNase A treatment) were resuspended in 200 µL of 10 mM Tris-HCl, pH 8.5.

### DNA isolation from bacteroids

*M. sativa* cv. Gabès and *M. truncatula* cv. A17 were used wild-type plants for all experiments. *M. truncatula dnf1*, *dnf2*, *dnf4*, *dnf5*, and *dnf7* mutants (Starker et al., 2006), derived from the A17 wild type, were used for collection of bacteroids blocked at various stages of differentiation. Seeds were scarified, surface sterilized, and germinated on Kalys agar as described previously (Kazmierczak et al., 2017). Fifty mL of overnight cultures of *E. meliloti* Rm2011 or FSM-MA, grown in YEB, were centrifuged (4,000 *g*, 20 minutes, room temperature) and resuspended in ~ 1,200 mL of sterile, distilled water to obtain a cell suspension at an OD_600nm_ of ~ 0.1. Germinated seedlings were immersed for one hour in the appropriate rhizobial cell suspension, and then planted in a perlite:sand (2:1) mixture. Plants were grown in a greenhouse for five to six weeks, with occasional watering with a 1 g L^−1^ nutrient solution (PlantProd solution [N-P-K, 0-15-40; Fertil]).

For whole nodule samples of wild type plants, pink nodules were collected from 53-60 plants per replicate 34-35 days post-inoculation; in the case of *dnf* mutants (and a matched wild-type A17 sample), nodules were collected from ~ 105 plants per genotype 23-24 days post-inoculation. Nodules were collected from the roots and stored in tubes in liquid nitrogen until collection was complete, at which point they were stored at −80°C until use. For sectioned nodule samples, pink nodules were collected from 103 *M. sativa* plants for each of the microsymbionts 35 to 40 days post-inoculation. Nodules were manually sectioned at the white-pink border. Nodule sections were stored in tubes over dry ice or liquid nitrogen until collection was complete, at which point they were stored at −80°C until use. Average plant shoot dry weights for all samples are listed in **Table S8**. Bacteroids were isolated from the nodule samples using Percoll gradient centrifugation as described elsewhere (Mergaert et al., 2006). The recovered bacteroids were resuspended in 50-100 µL of Bacteroid Extraction Buffer (BEB; 125 mM KCl, 50 mM Na-succinate, 50 mM TES, pH 7.0), and either used immediately for microscopy, flow cytometry, and DNA isolation, or stored at −80°C until use.

Nucleic acids were initially purified from most bacteroid samples using Epicentre MasterPure^TM^ Complete DNA and RNA Purification Kit, following the protocol for DNA isolation from cell samples; the exceptions were bacteroids purified from *dnf* mutant nodules (and the matched wild-type A17 sample), for which nucleic acids were isolated by using phenol:chloroform extractions followed by ammonium acetate DNA precipitations as described elsewhere (Cowie et al., 2006). For sectioned nodule samples, pure DNA was isolated by using the manufacturer’s protocol for the complete removal of RNA. For whole nodule samples, the isolated DNA was further purified by treating the nucleic acids samples with RNase A, after which pure DNA was isolated by using phenol:chloroform extractions followed by ammonium acetate DNA precipitations or alternatively using the MasterPure^TM^ DNA clean-up protocol for the DNA from *dnf* mutant nodules and the matched wild-type A17 sample. In all cases, the final DNA pellets were resuspended in 200 µL of 10 mM Tris-HCl, pH 8.5. Three biological replicates were performed for bacteroids isolated from most whole nodules, whereas only one replicate was performed for bacteroids isolated from sectioned nodules or *dnf* mutant nodules (and the matched wild-type A17 sample) due to low quantities of starting materials.

### DNA sequencing, modification detection, and motif analysis

DNA sequencing was performed at the U.S. Department of Energy Joint Genome Institute (JGI) or in-house at the University of Florence (the stationary phase samples and *dnf* mutant nodules and the matched wild-type A17 sample) using Pacific Biosciences (PacBio) sequencing technology (Eid et al., 2009). Genomic DNA was sheared to 3 kb using a Covaris LS220 (Covaris Inc., Woburn, MA, USA) or 15 kb (for stationary phase samples and bacteroids isolated from *dnf* mutant nodules and the matched wild-type A17 sample) using g-TUBEs (Covaris Inc., Woburn, MA, USA). Sheared DNA was treated with exonuclease to remove single-stranded ends and DNA damage repair mix followed by end repair and ligation of barcoded blunt adapters using SMRTbell Template Prep Kit 2.0 (PacBio, Menlo Park, CA, USA). Libraries were purified with AMPure PB beads (Beckman Coulter, Brea, CA, USA) and three or eight libraries with different barcodes were pooled at equimolar ratios and purified with AMPure PB beads. For most samples, SMRTbell template libraries were prepared using a Sequel Binding Kit 3.0 (PacBio, Menlo Park, CA, USA), and sequenced on a Sequel instrument using a v3 or v4 sequencing primer, 1M v3 SMRT cells, and Version 3.0 sequencing chemistry with 1×360 or 1×600 sequencing movie run times. The exceptions were the *E. meliloti* Rm2011 zone II and *E. meliloti* FSM-MA zone III bacteroid samples. For these samples, SMRTbell template libraries were prepared using a Sequel II Binding Kit 2.0 (PacBio, Menlo Park, CA, USA), and then sequenced on a Sequel II instrument using the tbd-sample dependent sequencing primer, 8M v1 SMRT cells, and Version 2.0 sequencing chemistry with 1×900 sequencing movie run times.

DNA modification detection and motif analysis were performed using the PacBio SMRT Link software (PacBio, Menlo Park, CA, USA). Briefly, raw reads were filtered using SFilter to remove short reads and reads derived from sequencing adapters. Filtered reads were aligned against the appropriate reference genome (**Table S2**) using BLASR (Chaisson and Tesler, 2012) and modified sites were then identified through kinetic analysis of the aligned DNA sequence data (Flusberg et al., 2010); the number of mapped bases per sample is provided in **Table S2**. Modified sites were then grouped into motifs using MotifFinder. These motifs represent the recognition sequences of MTase genes active in the genome (Clark et al., 2012). Downstream analyses were performed using custom Perl and R scripts.

### Flow cytometry

Flow cytometry was performed as described previously (Mergaert et al., 2006). Freshly prepared bacteroid samples were diluted in 200 µL of BEB, heat-treated for 10 minutes in a 70°C water bath, and then stained with the DNA-binding dye diamidino-2-phenylindole (DAPI). Cell size and ploidy level of the bacteroid samples were determined using flow cytometry with a Beckman Coulter CytoFLEX S instrument. Measurements consisted of 50,000 cells. Data analysis was performed using the CytExpert 2.2.0.97 software.

### Fluorescence microscopy

One µL of each freshly prepared bacteroid sample was mixed with 1 µL of 50 µg mL^−1^ DAPI or with both 1 µL of 50 µg mL^−1^ DAPI and 1 µL of 100 µg mL^−1^ propidium iodide (PI), which are both DNA binding dyes. Samples were visualized at 100x magnification under oil immersion using a Nikon Eclipse 80*i* fluorescence microscope with the NIS-Elements BR 4.00.01 software and a Digital Sight DS-U3 camera.

### Phylogenetic analysis

The nucleotide fasta files of representative *Ensifer* species were downloaded from the National Centre for Biotechnology Information (NCBI) Genome database. A core gene phylogeny was constructed using a previously prepared pipeline (diCenzo et al., 2018) reliant on the use of Roary 3.11.3 (Page et al., 2015), Prokka 1.12-beta (Seemann, 2014), PRANK 140110 (Löytynoja, 2014), trimAl (Capella-Gutiérrez et al., 2009), and RAxML 8.2.9 (Stamatakis, 2014). The phylogeny was visualized with the iTol webserver (Letunic and Bork, 2016). Identification of nodulation (*nodABC*) and nitrogenase genes (*nifHDK*) was performed with a published pipeline (diCenzo et al., 2018) reliant on the use of HMMER 3.1b2 (Eddy, 2009), and the Pfam-A 31.0 (Finn et al., 2016) and TIGERFAM 15.0 (Haft et al., 2013) databases.

## DATA AVAILABILITY

Most sequencing data is available through the JGI Genome Portal (genome.jgi.doe.gov/portal/) under Proposal 503835, as well as through NCBI (see **Table S2** for BioSample accessions). The data for stationary phase cultures and bacteroids isolated from *dnf* mutant nodules are available only through the NCBI (BioProject accessions PRJNA706182 and PRJNA705832; see **Table S2** for BioSample accessions). All custom scripts to perform the analyses described in this study are available through GitHub (github.com/diCenzo-Lab/003_2021_Ensifer_DNA_methylation), as are the flow cytometry FCS files.

## ACKNOWLEDGMENTS

We are grateful to E. Mullins (Teagasc, MTA2018233) for *E. adhaerens* OV14 and M. Bourge (I2BC) for help with the flow cytometry experiments.

