## Supplementary Tables and Figures for "DNA methylation in *Ensifer* species during free-living growth and during nitrogen-fixing symbiosis with *Medicago* spp."

**This PDF file includes:**

Figures S1 to S20
Tables S1 to S8
Legend for Datasets S1
SI References

**Other supplementary materials for this manuscript include the following:**

Datasets S1

**Table S1.** Putative methyltransferases in the *Sinorhizobium* pangenome and their distribution.

| Ortholog group | Annotation | Number of strains | Strains (locus tag) |
|---|---|---|---|
| 1 | Cell cycle-regulated methyltransferase CcrM | 20 | AK83 (Sinme_0634), B399 (BWO90_18765), B401 (BWO76_18415), BL225C (SinmeB_0539), FSM-MA (SMB554_04735), GR4 (C770_GR4Chr0926), HM006 (CDO22_06835), KH35c (CDO23_12830), KH46 (CDO24_07725), M162 (CDO25_05105), M270 (CDO26_11555), RMO17 (DU99_04775), RU11/001 (SMRU11_19915), Rm1021 (SMc00021), Rm41 (CDO27_16945), SM11 (SM11_chr0582), T073 (CDO28_05930), USDA1021 (CDO29_08105), USDA1106 (CDO30_03265), USDA1157 (CDO31_02955) |
| 2 | DNA (cytosine-5-)-methyltransferase | 2 | RU11/001 (SMRU11_31240), SM11 (SM11_chr0923) |
| 3 | Modification methylase | 1 | M270 (CDO26_10750) |
| 4 | DNA (cytosine-5-)-methyltransferase | 2 | USDA1106 (CDO30_09145), Rm1021 (SMc03763) |
| 5 | DNA (cytosine-5-)-methyltransferase | 1 | M270 (CDO26_06525) |
| 6 | Modification methylase | 1 | RU11/001 (SMRU11_01595) |
| 7 | DNA methyltransferase | 1 | T073 (CDO28_02045) |
| 8 | DNA modification methylase | 2 | RU11/001 (SMRU11_31210), SM11 (SM11_chr0917) |
| 9 | DNA methylase N-4/N-6 | 1 | AK83 (Sinme_2322) |
| 10 | Modification methylase | 1 | FSM-MA (SMB554_07065) |
| 11 | Site-specific DNA-methyltransferase | 1 | HM006 (CDO22_08570) |
| 12 | Site-specific DNA-methyltransferase | 1 | M270 (CDO26_04310) |
| 13 | DNA (cytosine-5-)-methyltransferase | 1 | RU11/001 (SMRU11_29360) |
| 14 | Site-specific DNA methylase | 1 | GR4 (C770_GR4pA023) |
| 15 | DNA cytosine methyltransferase | 1 | KH46 (CDO24_06020) |
| 16 | Modification methylase | 1 | KH46 (CDO24_13835) |
| 17 | DNA cytosine methyltransferase | 1 | HK46 (CDO24_34910) |
| 18 | Modification methylase | 1 | M270 (CDO26_04310) |
| 19 | DNA methylase | 1 | RU11/001 (SMRU11_01860) |
| 20 | SAM-dependent DNA methyltransferase | 1 | USDA1106 (CDO30_04960), Rm1021 (SMc02296) |
| 21 | N-6 DNA methylase | 1 | FSM-MA (SMB554_16155) |
| 22 | Type I restriction-modification system methyltransferase subunit | 1 | GR4 (C770_GR4Chr0590) |
| 23 | N-6 DNA methylase | 1 | KH35c (CDO23_13780) |
| 24 | SAM-dependent DNA methyltransferase | 1 | M162 (CDO25_03260) |

**Table S2.** Number of mapped subread bases per sample.

| Strain | Condition & Replicate | Genome Accession (NCBI Assembly) | Mapped Subreads Bases | NCBI BioSample Accessions for data generated in this study |
|---|---|---|---|---|
| *E. meliloti* FSM-MA | MM9-succinate 1 | GCA_002215195.1 | 2,414,122,786 | SAMN12793050 |
| *E. meliloti* FSM-MA | MM9-succinate 2 | GCA_002215195.1 | 1,973,904,434 | SAMN12793077 |
| *E. meliloti* FSM-MA | MM9-succinate 3 | GCA_002215195.1 | 2,276,105,908 | SAMN12793051 |
| *E. meliloti* FSM-MA | MM9-succinate stationary 1 | GCA_002215195.1 | 983,326,511 | SAMN18104108 |
| *E. meliloti* FSM-MA | MM9-succinate stationary 2 | GCA_002215195.1 | 1,149,234,588 | SAMN18104109 |
| *E. meliloti* FSM-MA | MM9-succinate stationary 3 | GCA_002215195.1 | 1,112,880,576 | SAMN18104110 |
| *E. meliloti* FSM-MA | *M. sativa* whole nodules 1 | GCA_002215195.1 | 669,928,620 | SAMN12793096 |
| *E. meliloti* FSM-MA | *M. sativa* whole nodules 2 | GCA_002215195.1 | 619,883,638 | SAMN12793097 |
| *E. meliloti* FSM-MA | *M. sativa* whole nodules 3 | GCA_002215195.1 | 2,014,796,418 | SAMN12792983 |
| *E. meliloti* FSM-MA | *M. truncatula* whole nodules 1 | GCA_002215195.1 | 1,223,643,922 | SAMN13167594 |
| *E. meliloti* FSM-MA | *M. truncatula* whole nodules 2 | GCA_002215195.1 | 993,061,044 | SAMN13168102 |
| *E. meliloti* FSM-MA | *M. truncatula* whole nodules 3 | GCA_002215195.1 | 1,038,438,145 | SAMN13167746 |
| *E. meliloti* FSM-MA | *M. sativa* distal nodule sections 1 | GCA_002215195.1 | 454,620,103 | SAMN15739021 |
| *E. meliloti* FSM-MA | *M. sativa* proximal nodule sections 1 | GCA_002215195.1 | 2,644,422,028 | SAMN14511029 |
| *E. meliloti* FSM-MA | *M. truncatula* A17 whole nodules 1 | GCA_002215195.1 | 900,330,949 | SAMN19249705 |
| *E. meliloti* FSM-MA | *M. truncatula dnf1* mutant whole nodules 1 | GCA_002215195.1 | 1,422,524,809 | SAMN19249700 |
| *E. meliloti* FSM-MA | *M. truncatula dnf2* mutant whole nodules 1 | GCA_002215195.1 | 668,179,310 | SAMN19249701 |
| *E. meliloti* FSM-MA | *M. truncatula dnf4* mutant whole nodules 1 | GCA_002215195.1 | 1,390,180,451 | SAMN19249702 |
| *E. meliloti* FSM-MA | *M. truncatula dnf5* mutant whole nodules 1 | GCA_002215195.1 | 3,414,132,613 | SAMN19249703 |
| *E. meliloti* FSM-MA | *M. truncatula dnf7* mutant whole nodules 1 | GCA_002215195.1 | 1,369,820,112 | SAMN19249704 |
| *E. meliloti* Rm2011 | MM9-succinate 1 | GCA_000346065.1 | 1,859,862,543 | SAMN12793063 |
| *E. meliloti* Rm2011 | MM9-succinate 2 | GCA_000346065.1 | 2,307,929,125 | SAMN12793100 |
| *E. meliloti* Rm2011 | MM9-succinate 3 | GCA_000346065.1 | 2,174,907,026 | SAMN12793065 |
| *E. meliloti* Rm2011 | MM9-succinate stationary 1 | GCA_000346065.1 | 1,303,384,501 | SAMN18104105 |
| *E. meliloti* Rm2011 | MM9-succinate stationary 2 | GCA_000346065.1 | 1,223,856,206 | SAMN18104106 |
| *E. meliloti* Rm2011 | MM9-succinate stationary 3 | GCA_000346065.1 | 1,217,560,804 | SAMN18104107 |
| *E. meliloti* Rm2011 | MM9-sucrose 1 | GCA_000346065.1 | 1,302,160,101 | SAMN12793017 |
| *E. meliloti* Rm2011 | MM9-sucrose 2 | GCA_000346065.1 | 1,673,395,654 | SAMN12793016 |
| *E. meliloti* Rm2011 | MM9-sucrose 3 | GCA_000346065.1 | 2,965,562,894 | SAMN12793099 |
| *E. meliloti* Rm2011 | *M. sativa* whole nodules 1 | GCA_000346065.1 | 814,031,492 | SAMN12793035 |
| *E. meliloti* Rm2011 | *M. sativa* whole nodules 2 | GCA_000346065.1 | 798,075,475 | SAMN12793034 |
| *E. meliloti* Rm2011 | *M. sativa* whole nodules 3 | GCA_000346065.1 | 1,138,170,578 | SAMN12793018 |
| *E. meliloti* Rm2011 | *M. sativa* distal nodule sections 1 | GCA_000346065.1 | 6,610,799,406 | SAMN16773451 |
| *E. meliloti* Rm2011 | *M. sativa* proximal nodule sections 2 | GCA_000346065.1 | 1,544,246,716 | SAMN13167882 |
| *E. meliloti* RmP3496 | MM9-sucrose 1 | GCA_000346065.1 | 1,138,116,503 | SAMN12792987 |
| *E. meliloti* RmP3496 | MM9-sucrose 2 | GCA_000346065.1 | 1,328,727,153 | SAMN12793078 |
| *E. meliloti* RmP3496 | MM9-sucrose 3 | GCA_000346065.1 | 2,234,156,208 | SAMN12793033 |

| | | | | |
|---|---|---|---|---|
| *E. fredii* NGR234 | MM9-succinate 1 | GCA_000018545.1 | 1,741,191,577 | SAMN12792974 |
| *E. fredii* NGR234 | MM9-succinate 2 | GCA_000018545.1 | 2,484,318,454 | SAMN12793102 |
| *E. fredii* NGR234 | MM9-succinate 3 | GCA_000018545.1 | 2,279,238,884 | SAMN12793103 |
| *E. adhaerens* OV14 | MM9-succinate 1 | GCA_000583045.1 | 1,883,494,303 | SAMN12793104 |
| *E. adhaerens* OV14 | MM9-succinate 2 | GCA_000583045.1 | 2,914,499,268 | SAMN12792975 |
| *E. adhaerens* OV14 | MM9-succinate 3 | GCA_000583045.1 | 2,272,958,147 | SAMN12793036 |

**Table S3.** Average extent of methylation of m6A modified motifs in *E. meliloti* Rm2011.

| Condition | Chromosome | pSymB | pSymA |
|---|---|---|---|
| GANTC | | | |
| Free-living (mid-exponential) | 0.878 | 0.916 | 0.883 |
| Free-living (stationary) | 0.957 | 0.960 | 0.964 |
| *M. sativa* distal nodule sections | 0.712 | 0.883 | 0.882 |
| *M. sativa* proximal nodule sections | 0.787 | 0.925 | 0.918 |
| *M. sativa* whole nodules | 0.791 | 0.921 | 0.920 |
| CGCA($N_5$)GTG | | | |
| Free-living (mid-exponential) | 0.978 | 0.980 | 0.981 |
| Free-living (stationary) | 0.974 | 0.981 | 0.982 |
| *M. sativa* distal nodule sections | 0.966 | 0.959 | 0.961 |
| *M. sativa* proximal nodule sections | 0.978 | 0.974 | 0.983 |
| *M. sativa* whole nodules | 0.973 | 0.972 | 0.976 |

**Table S4.** Average extent of methylation of m6A modified motifs in *E. meliloti* FSM-MA.

| Condition | Chromosome | pSymB | pSymA |
|---|---|---|---|
| GANTC | | | |
| Free-living (mid-exponential) | 0.803 | 0.898 | 0.836 |
| Free-living (stationary) | 0.938 | 0.947 | 0.947 |
| *M. sativa* distal nodule sections | 0.906 | 0.948 | 0.942 |
| *M. sativa* proximal nodule sections | 0.784 | 0.915 | 0.895 |
| *M. sativa* whole nodules | 0.860 | 0.946 | 0.936 |
| *M. truncatula* whole nodules | 0.793 | 0.917 | 0.911 |
| TCGA($N_8$)TCGA | | | |
| Free-living (mid-exponential) | 0.984 | 0.984 | 0.981 |
| Free-living (stationary) | 0.980 | 0.984 | 0.984 |
| *M. sativa* distal nodule sections | 0.973 | 0.973 | 0.980 |
| *M. sativa* proximal nodule sections | 0.978 | 0.915 | 0.977 |
| *M. sativa* whole nodules | 0.980 | 0.964 | 0.963 |
| *M. truncatula* whole nodules | 0.976 | 0.974 | 0.981 |

**Table S5.** Average extent of methylation of GANTC motifs in *E. meliloti* FSM-MA bacteroids purified from *M. truncatula dnf* mutant nodules and wild-type *M. truncatula* A17 nodules.

| *M. truncatula* genotype | Chromosome | pSymA | pSymB |
|---|---|---|---|
| GANTC | | | |
| *dnf1* | 0.859 | 0.868 | 0.874 |
| *dnf5* | 0.956 | 0.961 | 0.959 |
| *dnf2* | 0.870 | 0.939 | 0.941 |
| *dnf7* | 0.817 | 0.919 | 0.923 |
| *dnf4* | 0.814 | 0.915 | 0.924 |
| A17 | 0.804 | 0.917 | 0.920 |
| TCGA(N8)TCGA | | | |
| *dnf1* | 0.974 | 0.977 | 0.970 |
| *dnf5* | 0.984 | 0.985 | 0.983 |
| *dnf2* | 0.975 | 0.978 | 0.976 |
| *dnf7* | 0.974 | 0.972 | 0.974 |
| *dnf4* | 0.971 | 0.978 | 0.965 |
| A17 | 0.973 | 0.967 | 0.969 |

**Table S6.** Bacterial strains.

| Strain | Genotype | Source |
|---|---|---|
| *Ensifer adhearens* OV14 | Wild type OV14; not a nitrogen-fixing legume symbiont | (1) |
| *Ensifer fredii* NGR234 | Wild type NGR234 *rif-1*; $Rif^R$ | (2) |
| *Ensifer meliloti* FSM-MA | Wild type FSM-MA; $Cm^R$ | (3) |
| *Ensifer meliloti* Rm2011 | Wild type SU47 *str-3*; $Sm^R$ | Lab collection |
| *Ensifer meliloti* RmP3496 | Rm2011 lacking pSymA and pSymB; $Sm^R$ $Sp^R$ | (4, 5) |

Cm – Chloramphenicol; Sm – Streptomycin; Sp – Spectinomycin; Rif – Rifampicin

**Table S7.** Densities and doubling times of bacterial cultures grown for the isolation of DNA.

| | | Replicate 1 * | | | Replicate 2 | | | Replicate 3 | | |
|---|---|---|---|---|---|---|---|---|---|---|
| Strain | Carbon source | Starting density † | Final density | Doubling time (h) | Starting density | Final density | Doubling time (h) | Starting density | Final density | Doubling time (h) |
| *E. meliloti* Rm2011 | Succinate | 0.015 | 0.554 | 2.98 | 0.014 | 0.525 | 2.94 | 0.014 | 0.499 | 2.98 |
| *E. meliloti* FSM-MA | Succinate | 0.004 | 0.571 | 2.13 | 0.003 | 0.420 | 2.20 | 0.003 | 0.369 | 2.26 |
| *E. fredii* NGR234 | Succinate | 0.019 | 0.631 | 3.09 | 0.016 | 0.650 | 2.92 | 0.016 | 0.634 | 2.94 |
| *E. adhaerens* OV14 | Succinate | 0.002 | 0.686 | 1.86 | 0.002 | 0.564 | 1.84 | 0.002 | 0.552 | 1.85 |
| *E. meliloti* Rm2011 | Sucrose | 0.014 | 0.581 | 2.91 | 0.013 | 0.512 | 2.90 | 0.013 | 0.516 | 2.90 |
| *E. meliloti* RmP3496 | Sucrose | 0.039 | 0.574 | 3.98 | 0.035 | 0.389 | 4.44 | 0.035 | 0.401 | 4.38 |

* Each Replicate 1 sample consisted of two 40 mL cultures (inoculated from the same starter culture) that were combined prior to DNA isolation. The density and doubling time values for this replicate are based on just one of the cultures for each sample.

† All density values are $OD_{600nm}$ measurements.

**Table S8.** Plant shoot dry weights.

| Plant | Bacterium | Replicate | Number of plants | Average shoot dry weight (mg) |
|---|---|---|---|---|
| Plants for whole nodules † | | | | |
| *M. sativa* | *E. meliloti* Rm2011 | 1 | 55 | 57 |
| *M. sativa* | *E. meliloti* Rm2011 | 2 | 52 | 60 |
| *M. sativa* | *E. meliloti* Rm2011 | 3 | 52 | 54 |
| *M. sativa* | *E. meliloti* FSM-MA | 1 | 60 | 84 |
| *M. sativa* | *E. meliloti* FSM-MA | 2 | 60 | 80 |
| *M. sativa* | *E. meliloti* FSM-MA | 3 | 55 | 90 |
| *M. truncatula* | *E. meliloti* FSM-MA | 1 | 60 | 58 |
| *M. truncatula* | *E. meliloti* FSM-MA | 2 | 60 | 60 |
| *M. truncatula* | *E. meliloti* FSM-MA | 3 | 55 | 68 |
| *M. sativa* | Uninoculated | 1 | 10 | ND * |
| *M. truncatula* | Uninoculated | 1 | 5 | ND * |
| Plants for sectioned nodules† | | | | |
| *M. sativa* | *E. meliloti* Rm2011 | 1 | 103 | 180 |
| *M. sativa* | *E. meliloti* FSM-MA | 1 | 103 | 162 |
| *M. sativa* | Uninoculated | 1 | 55 | 31 |

* ND: Not determined.

† Plants for isolation of whole nodules and plants for isolation of sectioned nodules were grown independently at separate periods of the year.

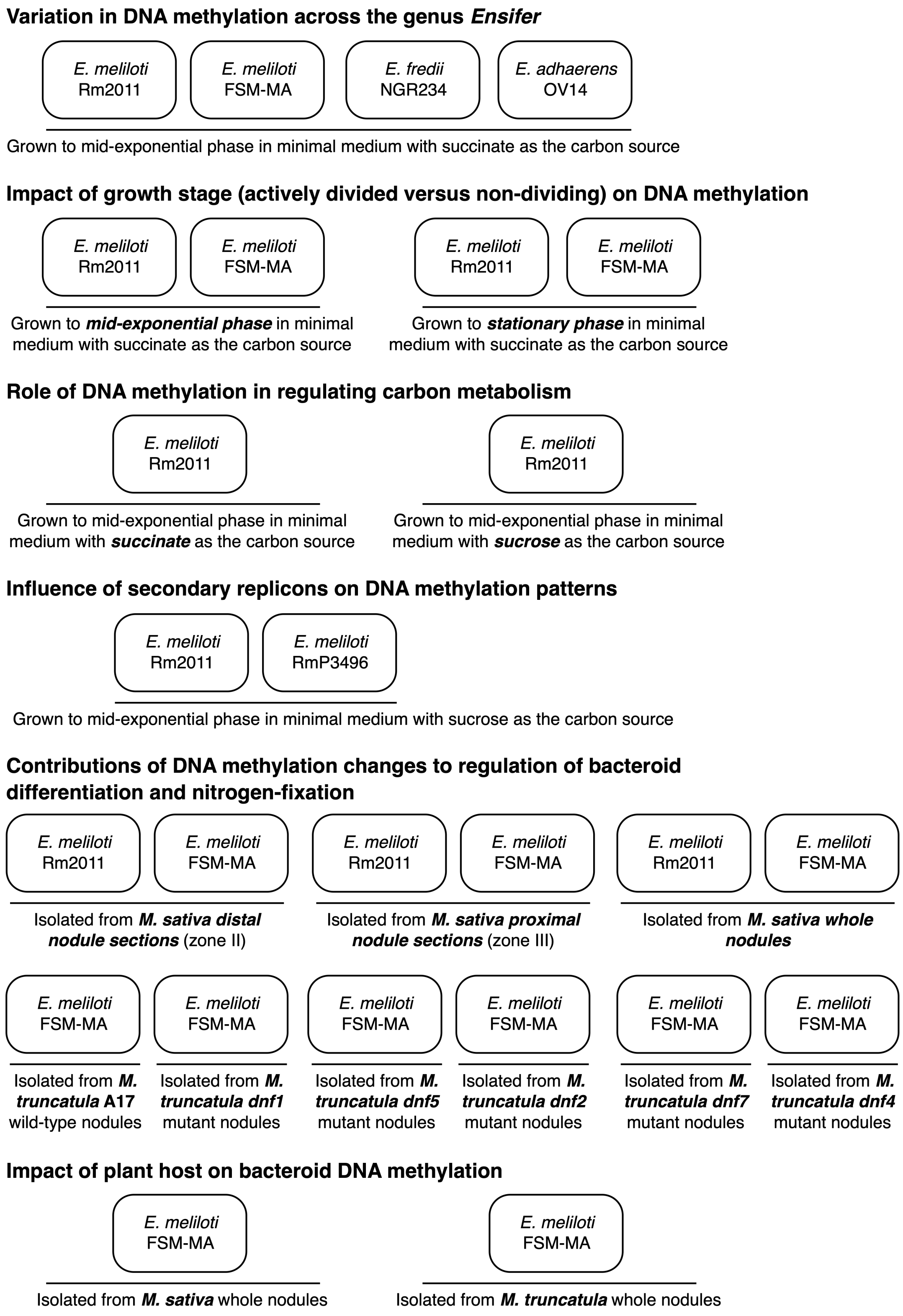


**Figure S1. Experimental design overview.** A schematic overview of the experimental design of this study, summarizing the comparisons that were performed, and which strains/conditions correspond to each comparison.

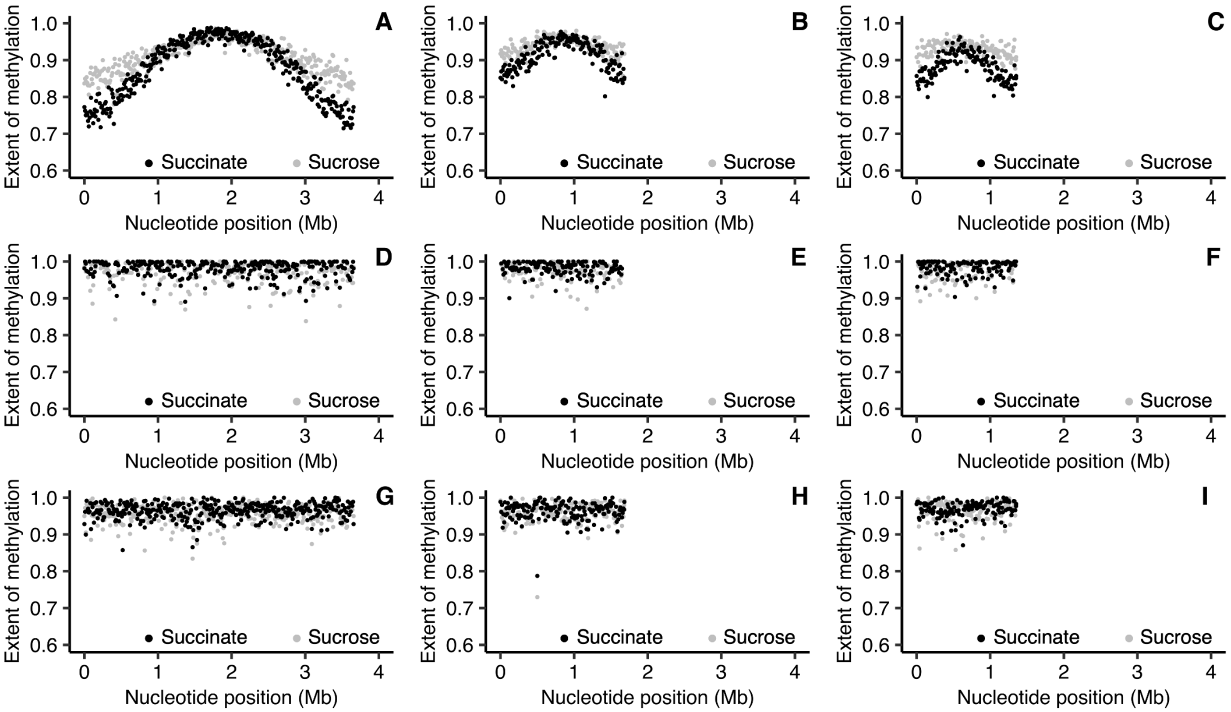


**Figure S2. Impact of carbon source on genome-wide DNA methylation of *E. meliloti* Rm2011.** The extent of methylation is shown, using a 10 kb sliding window, of cells grown to mid-exponential phase and provided succinate (black) or sucrose (grey) as the sole source of carbon. (**A**-**C**) Data for the GANTC motif for the chromosome (A), pSymB (B), and pSymA (C). (**D**-**F**) Data for the CGCA($N_5$)GTG motif for the chromosome (D), pSymB (E), and pSymA (F). (**G**-**I**) Data for the RCGCCTC motif for the chromosome (G), pSymB (H)**,** and pSymA (I).

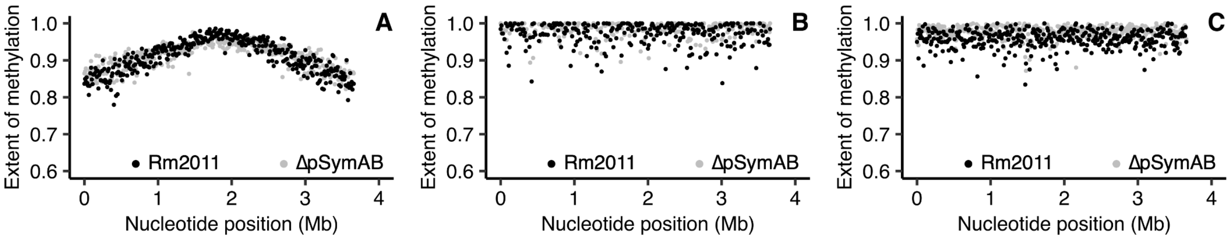


**Figure S3. Impact of pSymA and pSymB removal on chromosome-wide DNA methylation of *E. meliloti* Rm2011.** The extent of methylation for the chromosome, shown using a 10 kb sliding window, is provided for wild type *E. meliloti* Rm2011 (black) or *E. meliloti* ΔpSymAB (grey) grown to mid-exponential phase and provided sucrose as the sole source of carbon. (**A**) Data for the GANTC motif. (**B**) Data for the CGCA($N_5$)GTG motif. (**C**) Data for the RCGCCTC motif.

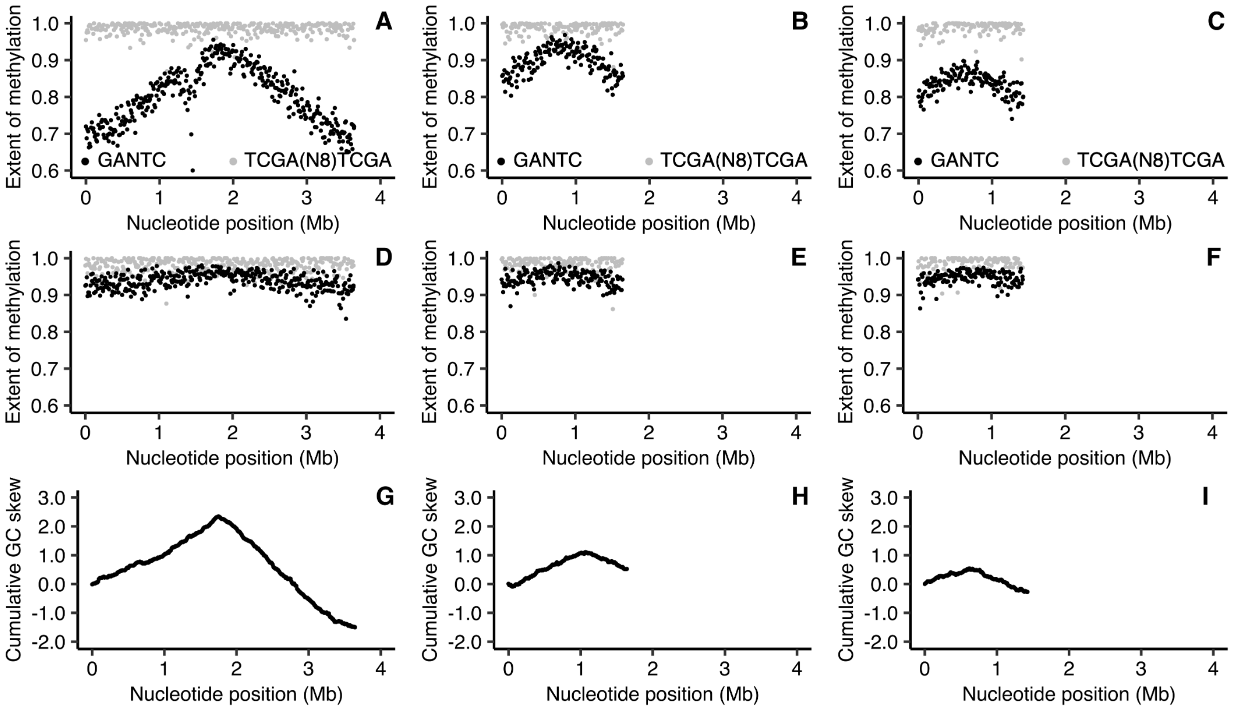


**Figure S4. Genome-wide DNA methylation of *E. meliloti* FSM-MA.** (**A**-**F**) The extent of methylation is shown, using a 10 kb sliding window, of GANTC sites (black) and TCGA($N_5$)TCGA sites (grey) across the chromosome (**A**,**D**), pSymB (**B**,**E**), and pSymA (**C**,**F**) replicons of exponential phase (**A**-**C**) or early stationary phase (**D**-**F**) *E. meliloti* FSM-MA. Averages from three biological replicates are shown. (**G**-**I**) Cumulative GC skews are shown, using a 10 kb sliding window, across the *E. meliloti* FSM-MA chromosome (**G**), pSymB (**H**), and pSymA (**I**) replicons.

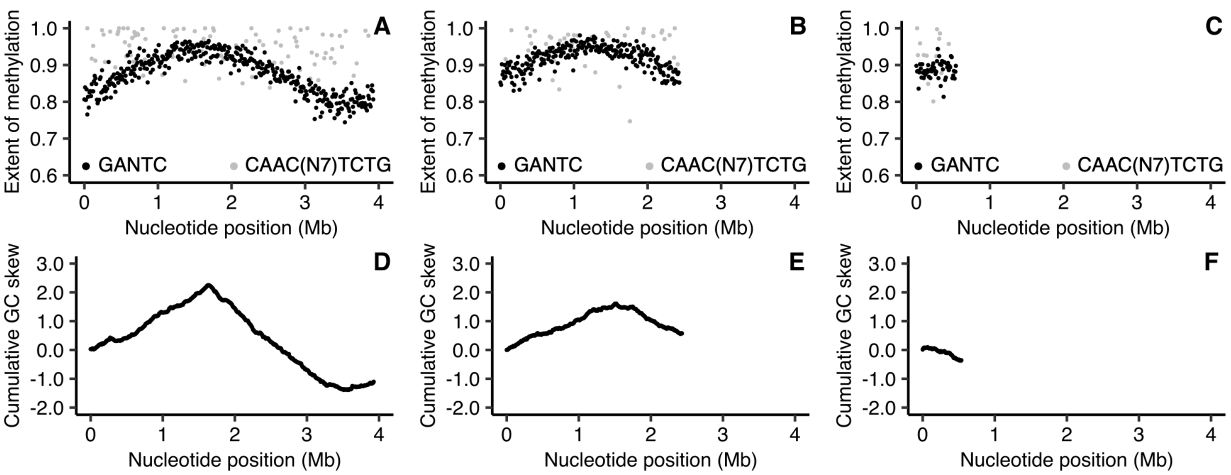


**Figure S5. Genome-wide DNA methylation of *E. fredii* NGR234.** (**A**-**C**) The extent of methylation is shown, using a 10 kb sliding window, of GANTC sites (black) and CAAC($N_7$)TCTG sites (grey) across the chromosome (**A**), pNGR234b (**B**), and pNGR234a (**C**) replicons of exponential phase *E. fredii* NGR234. Averages from three biological replicates are shown. (**D**-**F**) Cumulative GC skews are shown, using a 10 kb sliding window, across the *E. fredii* NGR234 chromosome (**D**), pNGR234b (**E**), and pNGR234a (**F**) replicons. Nucleotide positions refer to the nucleotide positions in the genome assembly files available through NCBI, which does not necessarily correlate with the location of the origin of replication. Instead, the origin of replication and replication terminus regions of each replicon are expected to be represented by the low and high points, respectively, on the cumulative GC skews.

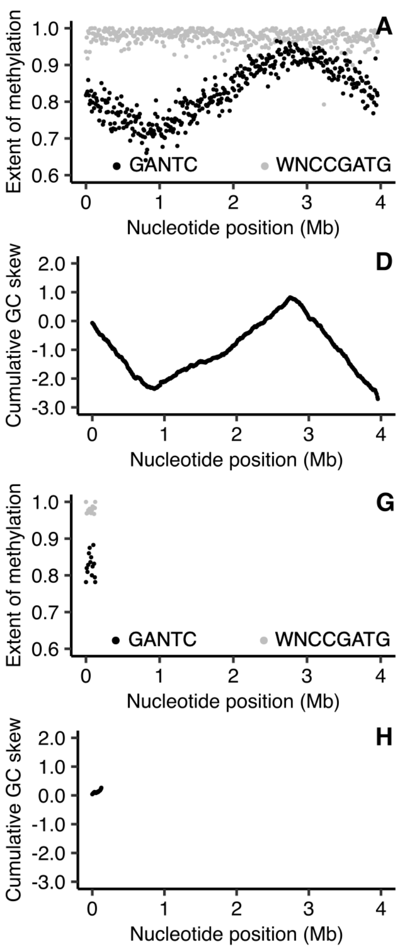


**Figure S6. Genome-wide DNA methylation of *E. adhaerens* OV14.** (**A**-**C**,**G**) The extent of methylation is shown using a 10 kb sliding window, of GANTC sites (black) and WNCCGATG sites (grey) across chromosome 1 (**A**), chromosome 2 (**B**), pOV14b (**C**), and pOV14c (**G**) replicons of exponential phase *E. adhaerens* OV14. Averages from three biological replicates are shown. (**D**-**F**,**H**) Cumulative GC skews are shown, using a 10 kb sliding window, across the *E. adhaerens* OV14 chromosome 1 (**D**), chromosome 2 (**E**), pOV14b (**F**), and pOV14c (**H**) replicons. Nucleotide positions refer to the nucleotide positions in the genome assembly files available through NCBI, which does not necessarily correlate with the location of the origin of replication. Instead, the origin of replication and replication terminus regions of each replicon are expected to be represented by the low and high points, respectively, on the cumulative GC skews.

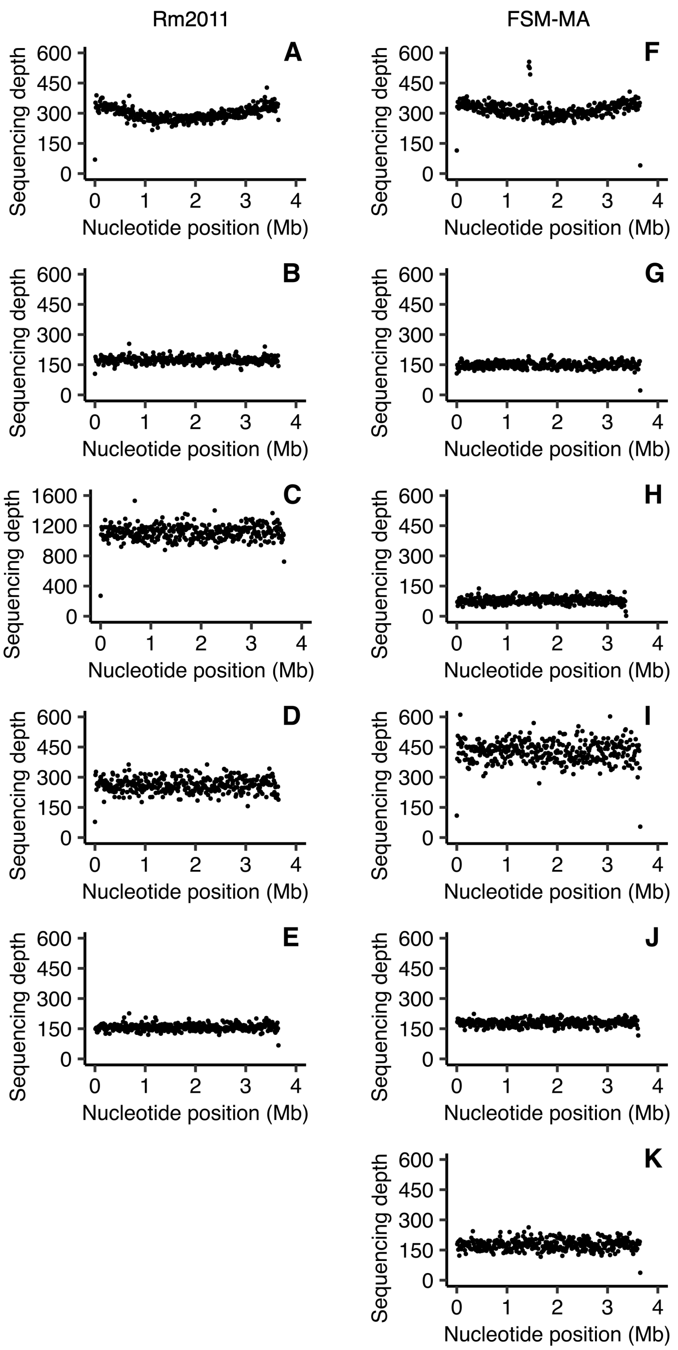


**Figure S7. Sequencing depth across the *E. meliloti* RM2011 and FSM-MA chromosome.** The sequencing depths (i.e., number of reads mapping to a given nucleotide) of the (**A**-**E**) *E. meliloti* Rm2011 chromosome and (**F**-**K**) *E. meliloti* FSM-MA chromosome are shown, using a 10 kb sliding window. Averages from three biological replicates are shown for free-living and whole nodule samples; data represents one replicate for the zone II and zone III nodule sections. (**A**,**F**) Free-living cells harvested in mid-exponential phase. (**B**,**G**) Free-living cells harvested in early stationary phase. (**C**,**H**) Bacteroids isolated from *M. sativa* zone II nodule sections. (**D**,**I**) Bacteroids isolated from *M. sativa* zone III nodule sections. (**E**,**J**) Bacteroids isolated from *M. sativa* whole nodule samples. (**K**) Bacteroids isolated from *M. truncatula* whole nodule samples.

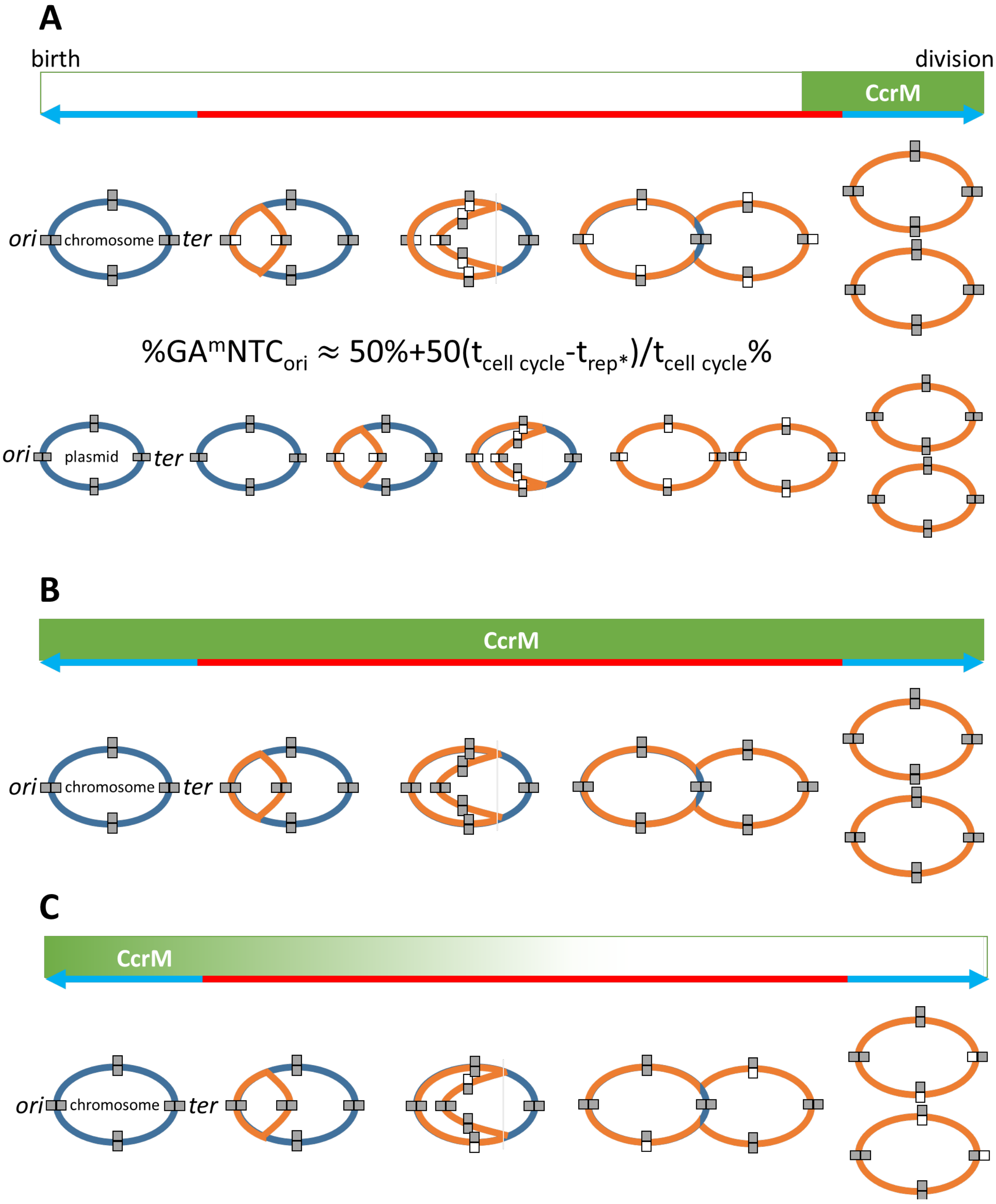


**Figure S8. Model describing the methylation patterns observed in cultures and bacteroids.** **(A)** Methylation pattern in free-living cells in exponential and stationary phases of growth. The blue/red line indicates the relevant cell cycle phases: red is the S-phase or genome replication phase; blue is the gap phases and division. The replication progression and methylation status of the circular chromosome and a plasmid is depicted during the cell cycle progression from cell birth until division. The replicons at cell birth are in dark blue and the newly replicated DNA is in orange. Full grey rectangles indicate the fully methylated state and grey/white rectangles indicate the hemimethylated state. Replicons start the cell cycle in a fully methylated state across the genome and freshly replicated genome segments become hemimethylated. CcrM activity (green box) is confined to a short window in the cell cycle because the *ccrM* gene is expressed in the late phase of genome replication (6) and because the protein is degraded by the Lon protease prior to cell division (7). The formula expresses the extent of methylation at the origin (*ori*) of replication in an asynchronous bacterial population in culture. The methylation level shuttles between 50% and 100% depending on the proportion of cells in the culture that have a hemimethylated origin due to ongoing DNA replication prior to the activation of CcrM, and the complementary proportion of cells that have a fully methylated origin because they either have not yet started DNA replication or they have completed (or nearly completed) DNA replication and have activated CcrM. The fraction of cells in the different cell phases is in turn proportional to the duration (t) of the corresponding cell cycle stages ($t_{rep*}$ is the time from initiation of replication till activation of CcrM, just before replication termination). Based on this formula, our experimentally measured extent of methylation at the *ori* of the chromosome in exponential phase cells, and the measured doubling time of *E. meliloti*, we estimate that it takes about 1 hour to replicate the chromosome, corresponding to a DNA polymerase rate of ~ 450 to 500 nucleotides per second. This compares well with the estimated DNA polymerase rates of *Escherichia coli* and *Caulobacter crescentus*, which are ~ 600 and ~ 350 nucleotides per second, respectively (8, 9). In the genus *Ensifer*, the extent of methylation at the *ori* of the megaplasmids is higher than in the chromosome, while the terminus (*ter*) of the megaplasmids has a slightly lower extent of methylation. Since DNA methylation happens at the fixed stage of the cell cycle at the end of chromosome replication, it follows that plasmid replication is initiated later in the cell cycle than initiation of chromosome replication, and that their replication terminates slightly before termination of the chromosome and the activation of CcrM. This is consistent with independent analyses revealing spatiotemporal regulation of DNA replication and partitioning in *E. meliloti* (6, 10). The *ter* of the chromosome remains near fully methylated during the complete cell cycle because its replication coincides with the activity window of CcrM. In a stationary phase culture, methylation approaches 100% because few cells are in the replication phase and the majority of cells are in a gap phase of the cell cycle. **(B)** Methylation pattern in an early stage of bacteroid differentiation. Early stage bacteroids have a high extent of methylation across the genome suggesting that the activity of CcrM is extended to include also the replication phase of the cell cycle. This can be the result of an aberrant expression of the *ccrM* gene or a lack of proteolytic degradation of the CcrM protein. **(C)** Methylation pattern in a late stage of bacteroid differentiation. In mature bacteroids, the methylation extent at the *ori* is high but characteristically, the *ter* of the chromosome in bacteroids has a reduced methylation extent. This can be the result of a drop in CcrM activity during the last chromosome replication cycle of the endoreduplication process in differentiating bacteroids. Possibly, the drop of CcrM activity can also be the cause of the arrest of the endoreduplication process. Reduced methylation at the *ter* region is generally not observed in the plasmids, suggesting that plasmid endoreduplication finishes before chromosome endoreduplication and that CcrM activity stops after the end of plasmid replication. The iconography of the illustrations is based on Figure 1 of Mohapatra et al. 2014 (11).

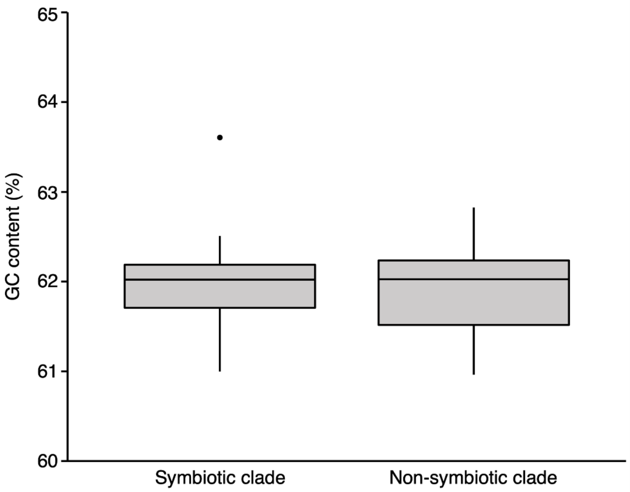


**Figure S9. GC content in the genus *Ensifer*.** Box plots summarizing the GC content of the genomes of 157 *Ensifer* strains are shown. The monophyletic "symbiotic" and "non-symbiotic" clades as defined previously (12), are represented by 111 and 44 genomes respectively.

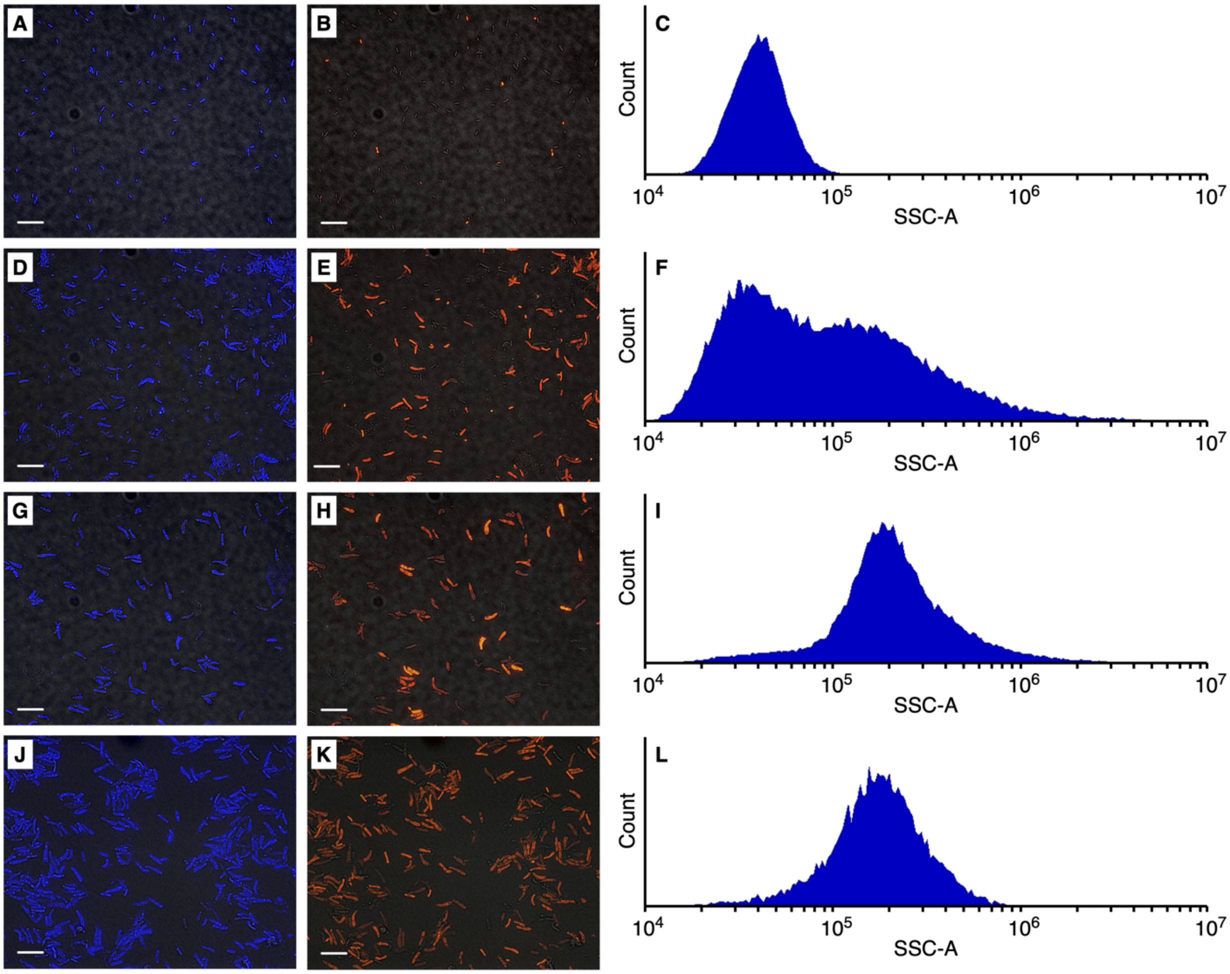


**Figure S10. Morphology of *E. meliloti* Rm2011 cell populations.** Free-living *E. meliloti* Rm2011 cells (**A**-**C**), and *E. meliloti* Rm2011 cells purified from *M. sativa* zone II nodule sections (**D**-**F**), *M. sativa* zone III nodule sections (**G**-**I**), or *M. sativa* whole nodules (**J**-**L**). Ploidy data is provided in Figure S11. Micrographs show *E. meliloti* Rm2011 cell populations stained with the DNA binding dyes DAPI (blue) and PI (red). The scale bar represents 10 µm. (**A**,**D**,**G**,**J**) DAPI fluorescence overlaid with DIC (differential interference contrast) images. (**B**,**E**,**H**,**K**) PI fluorescence overlaid with DIC images. (**C**,**F**,**I**,**L**) Histograms summarizing the distribution of flow cytometry side scattering values, providing an estimation of cell morphology, of heat-killed *E. meliloti* Rm2011 populations. Graphs are based on 50,000 cells.

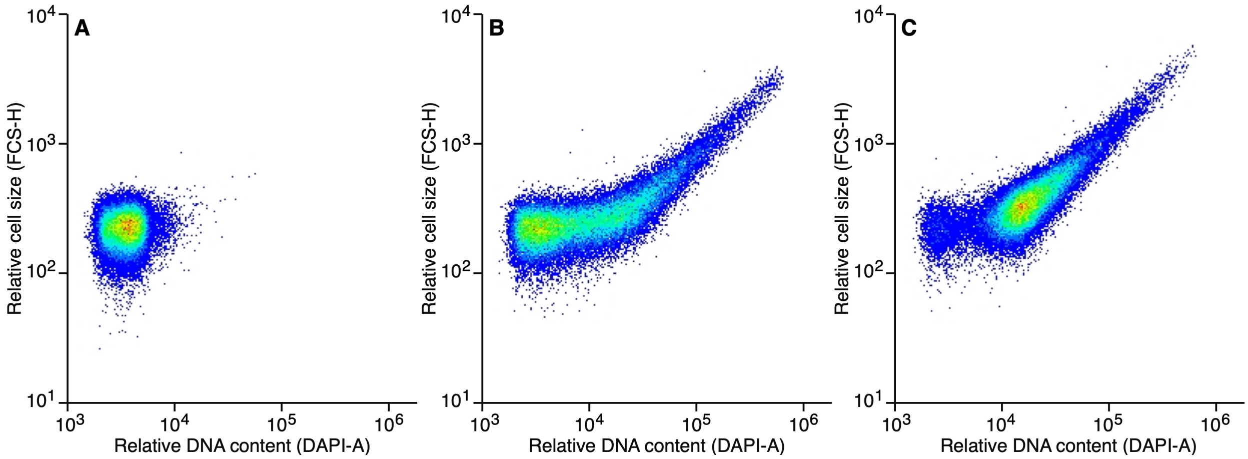


**Figure S11. Cell size and DNA content of *E. meliloti* Rm2011 cell populations.** Pseudo-coloured scatterplots displaying the DNA content (X-axis) and cell size (Y-axis) of various *E. meliloti* Rm2011 populations, based on flow cytometry readings of 50,000 cells. The colour scheme indicates the number of values plotted at a given location of the graph, with blue to red indicating lower to higher density. Free-living *E. meliloti* Rm2011 cells (**A**), *E. meliloti* Rm2011 cells purified from *M. sativa* zone II nodule sections (**B**), and *E. meliloti* Rm2011 cells purified from *M. sativa* zone III nodule sections (**C**).

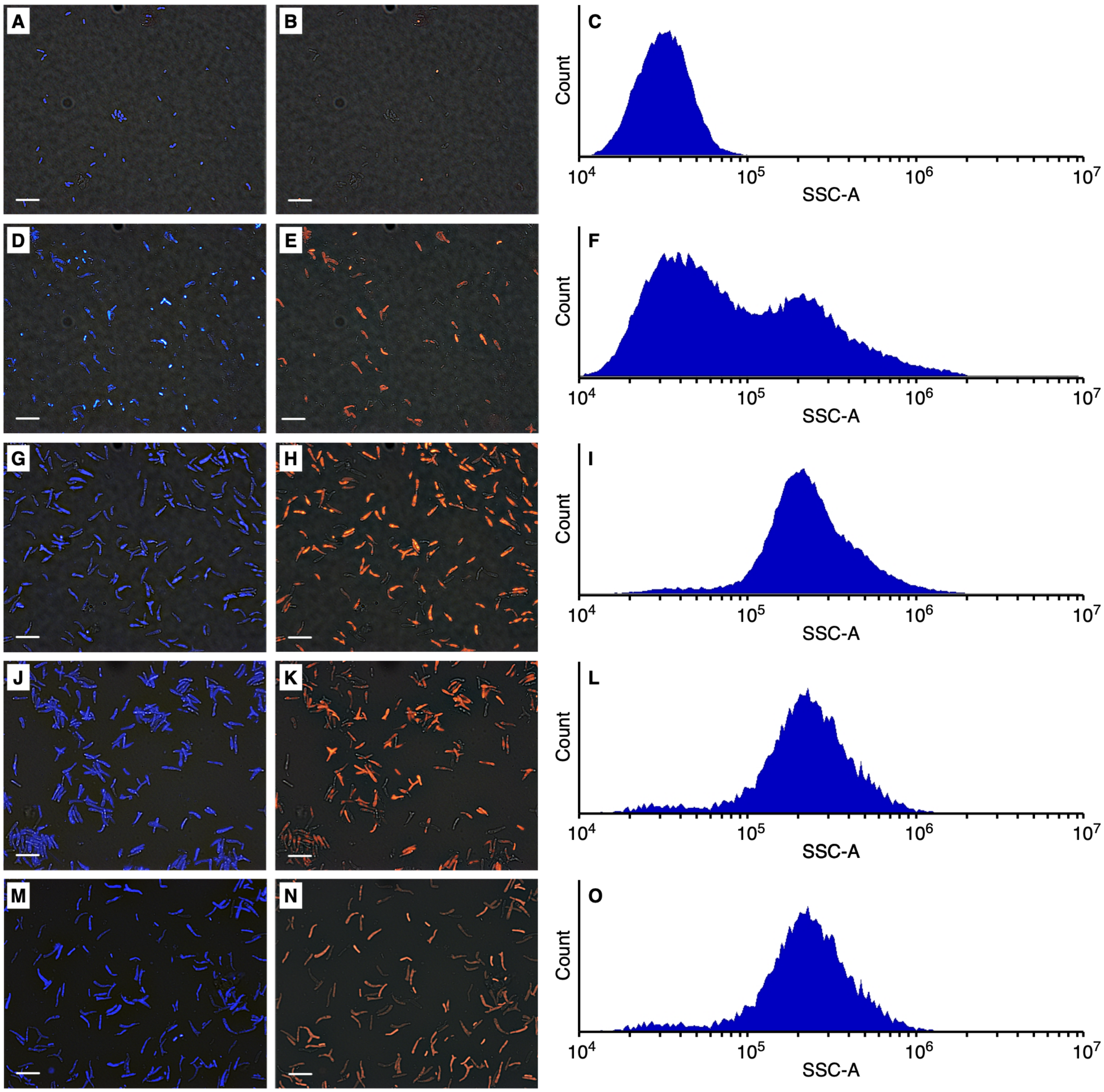


**Figure S12. Morphology of *E. meliloti* FSM-MA cell populations.** Free-living *E. meliloti* FSM-MA cells (**A**-**C**), and *E. meliloti* FSM-MA cells purified from *M. sativa* zone II nodule sections (**D**-**F**), *M. sativa* zone III nodule sections (**G**-**I**), *M. sativa* whole nodules (**J**-**L**), or *M. truncatula* whole nodules (**M-O**). Ploidy data is provided in Figure S13. Micrographs show *E. meliloti* FSM-MA cell populations stained with the DNA binding dyes DAPI (blue) and PI (red). The scale bar represents 10 µm. (**A**,**D**,**G**,**J**,**M**) DAPI fluorescence overlaid with DIC (differential interference contrast) images. (**B**,**E**,**H**,**K**,**N**) PI fluorescence overlaid with DIC images. (**C**,**F**,**I**,**L**,**O**) Histograms summarizing the distribution of flow cytometry side scattering values of heat-killed *E. meliloti* FSM-MA populations. Graphs are based on 50,000 cells.

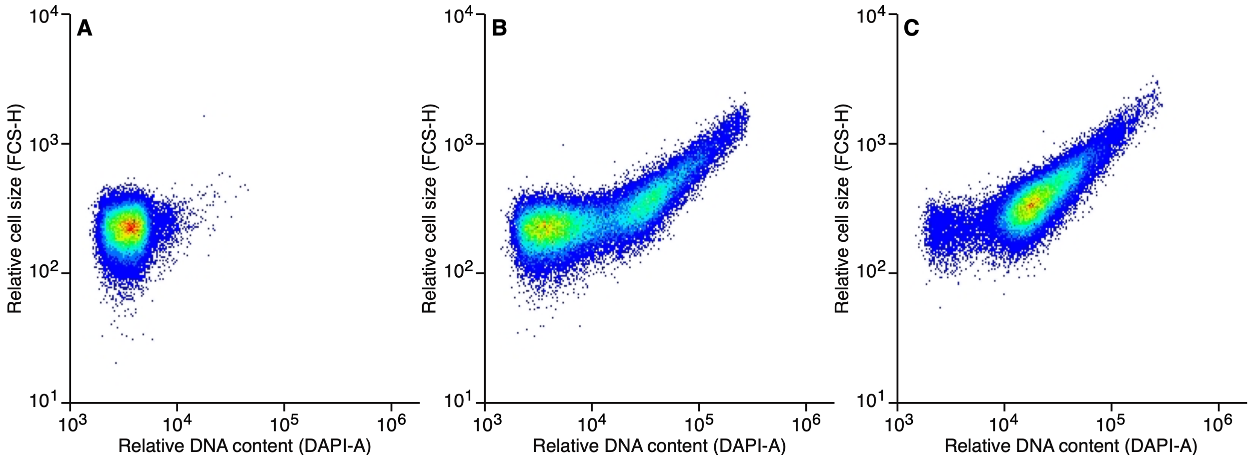


**Figure S13. Cell size and DNA content of *E. meliloti* FSM-MA cell populations.** Pseudo-coloured scatterplots displaying the DNA content (X-axis) and size (Y-axis) of various *E. meliloti* FSM-MA populations, based on flow cytometry readings of 50,000 cells. The colour scheme indicates the number of values plotted at a given location of the graph, with blue to red indicating lower to higher density. Free-living *E. meliloti* FSM-MA cells (**A**), *E. meliloti* FSM-MA cells purified from *M. sativa* zone II nodule sections (**B**), and *E. meliloti* FSM-MA cells purified from *M. sativa* zone III nodule sections (**C**).

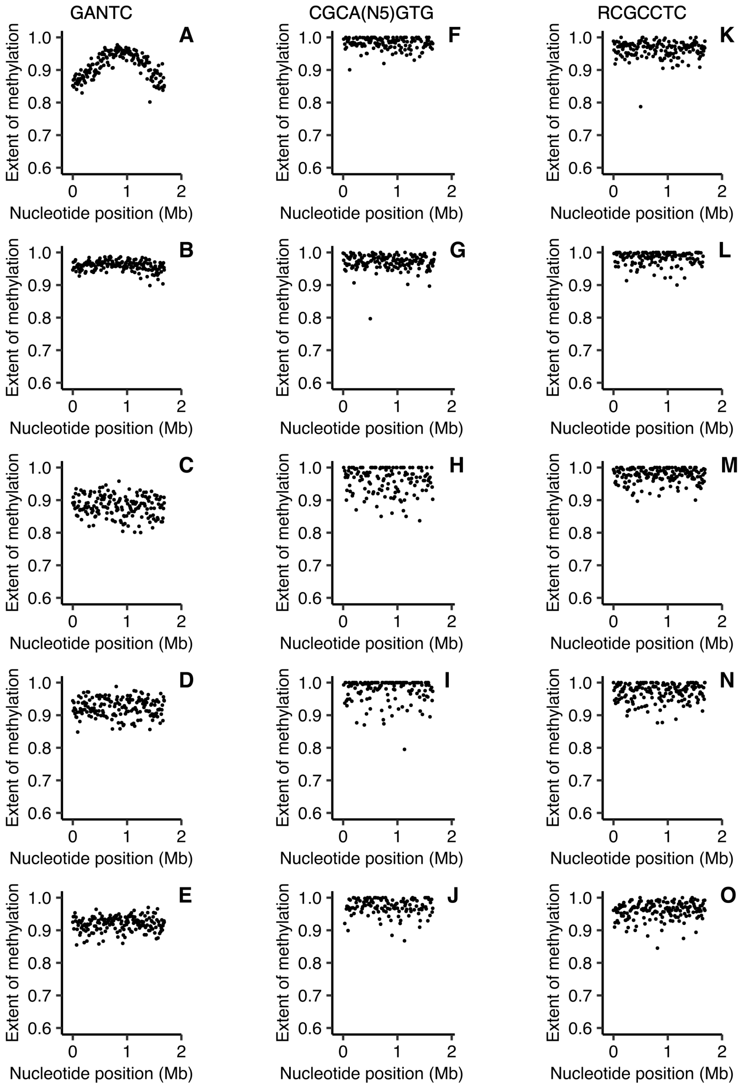


**Figure S14. pSymB-wide DNA methylation of *E. meliloti* Rm2011 bacteroids.** The extent of methylation of (**A**-**E**) GANTC, (**F**-**J**) CGCA($N_5$)GTG, and (**K**-**O**) RCGCCTC motifs across the *E. meliloti* Rm2011 pSymB replicon is shown using a 10 kb sliding window. Averages from three biological replicates are shown for free-living and whole nodule samples; data represents one replicate for the zone II and zone III nodule sections. (**A**,**F**,**K**) Free-living cells harvested in mid-exponential phase. (**B**,**G**,**L**) Free-living cells harvested in early stationary phase. (**C**,**H**,**M**) Bacteroids isolated from *M. sativa* zone II nodule sections. (**D**,**I**,**N**) Bacteroids isolated from *M. sativa* zone III nodule sections. (**E**,**J**,**O**) Bacteroids isolated from *M. sativa* whole nodule samples.

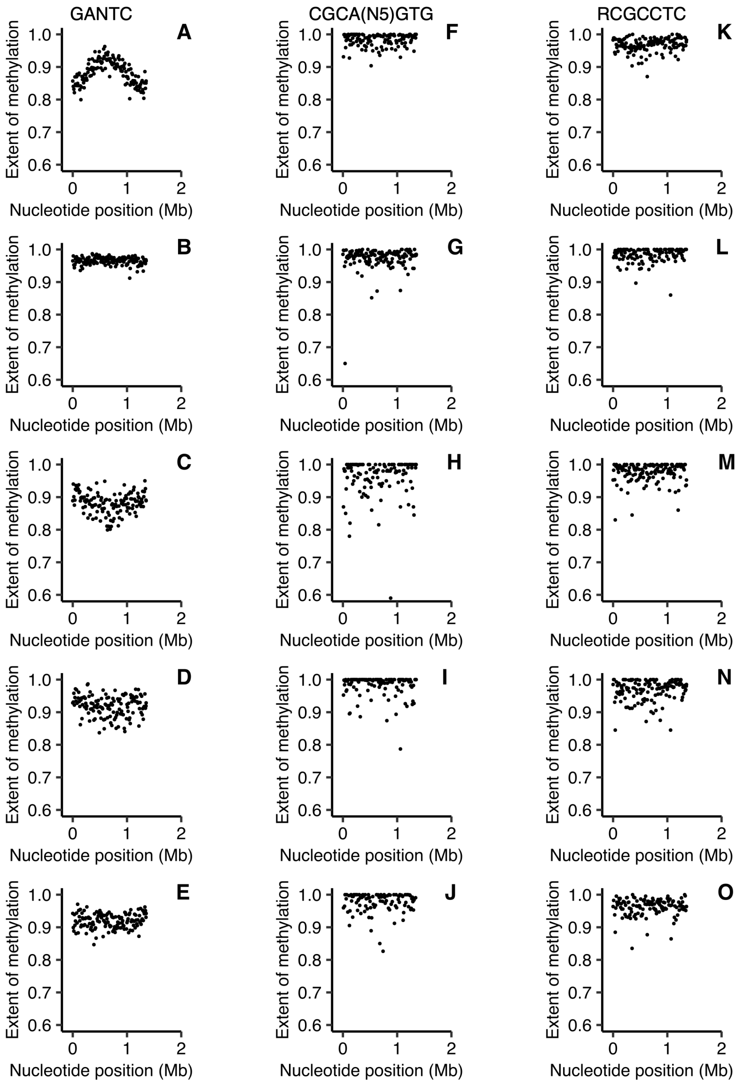


**Figure S15. pSymA-wide DNA methylation of *E. meliloti* Rm2011 bacteroids.** The extent of methylation of (**A**-**E**) GANTC, (**F**-**J**) CGCA($N_5$)GTG, and (**K**-**O**) RCGCCTC motifs across the *E. meliloti* Rm2011 pSymA replicon is shown using a 10 kb sliding window. Averages from three biological replicates are shown for free-living and whole nodule samples; data represents one replicate for the zone II and zone III nodule sections. (**A**,**F**,**K**) Free-living cells harvested in mid-exponential phase. (**B**,**G**,**L**) Free-living cells harvested in early stationary phase. (**C**,**H**,**M**) Bacteroids isolated from *M. sativa* zone II nodule sections. (**D**,**I**,**N**) Bacteroids isolated from *M. sativa* zone III nodule sections. (**E**,**J**,**O**) Bacteroids isolated from *M. sativa* whole nodule samples.

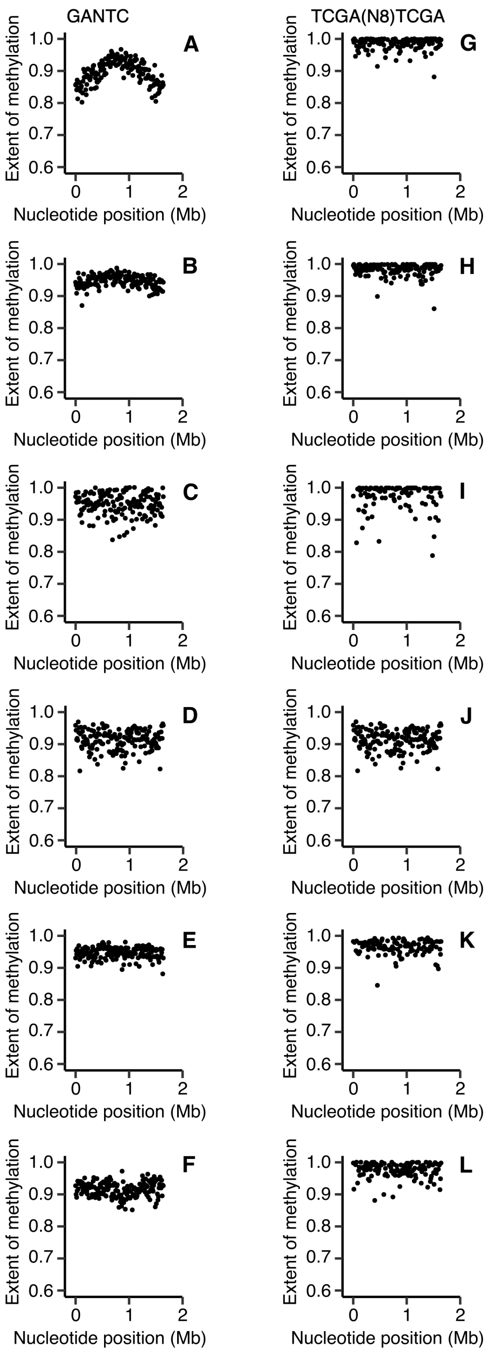


**Figure S16. pSymB-wide DNA methylation of *E. meliloti* FSM-MA bacteroids.** The extent of methylation of (**A**-**F**) GANTC and (**G**-**L**) TCGA($N_8$)TCGA motifs across the *E. meliloti* FSM-MA pSymB replicon is shown using a 10 kb sliding window. Averages from three biological replicates are shown for free-living and whole nodule samples; data represents one replicate for the zone II and zone III nodule sections. (**A**,**G**) Free-living cells harvested in mid-exponential phase. (**B**,**H**) Free-living cells harvested in early stationary phase. (**C**,**I**) Bacteroids isolated from *M. sativa* zone II nodule sections. (**D**,**J**) Bacteroids isolated from *M. sativa* zone III nodule sections. (**E**,**K**) Bacteroids isolated from *M. sativa* whole nodule samples. (**F**,**L**) Bacteroids isolated from *M. truncatula* whole nodule samples.

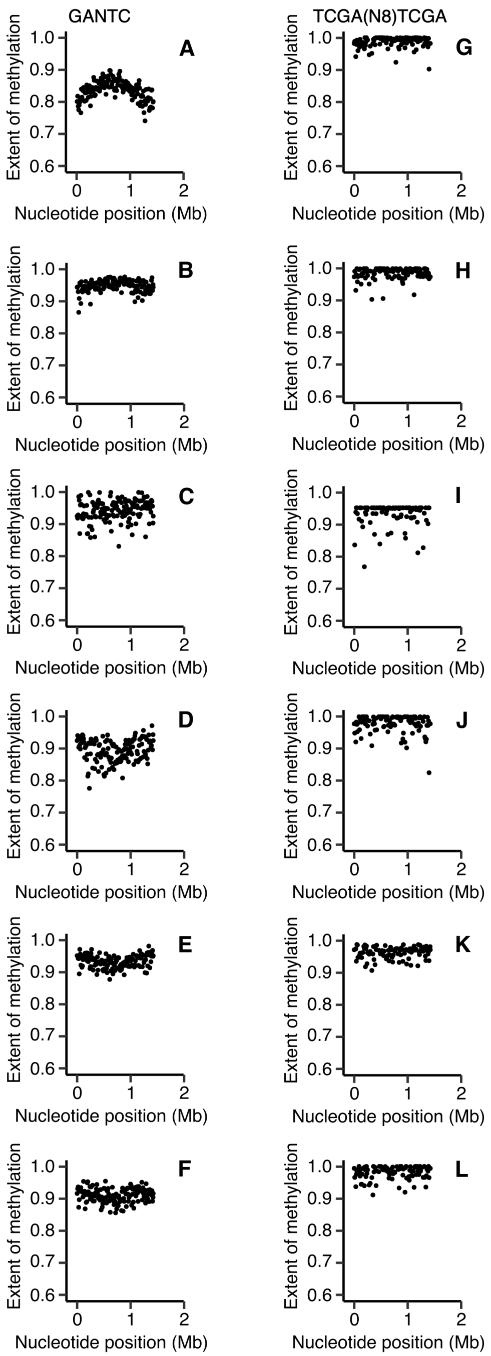


**Figure S17. pSymA-wide DNA methylation of *E. meliloti* FSM-MA bacteroids.** The extent of methylation of (**A**-**F**) GANTC and (**G**-**L**) TCGA($N_8$)TCGA motifs across the *E. meliloti* FSM-MA pSymA replicon is shown using a 10 kb sliding window. Averages from three biological replicates are shown for free-living and whole nodule samples; data represents one replicate for the zone II and zone III nodule sections. (**A**,**G**) Free-living cells harvested in mid-exponential phase. (**B**,**H**) Free-living cells harvested in early stationary phase. (**C**,**I**) Bacteroids isolated from *M. sativa* zone II nodule sections. (**D**,**J**) Bacteroids isolated from *M. sativa* zone III nodule sections. (**E**,**K**) Bacteroids isolated from *M. sativa* whole nodule samples. (**F**,**L**) Bacteroids isolated from *M. truncatula* whole nodule samples.

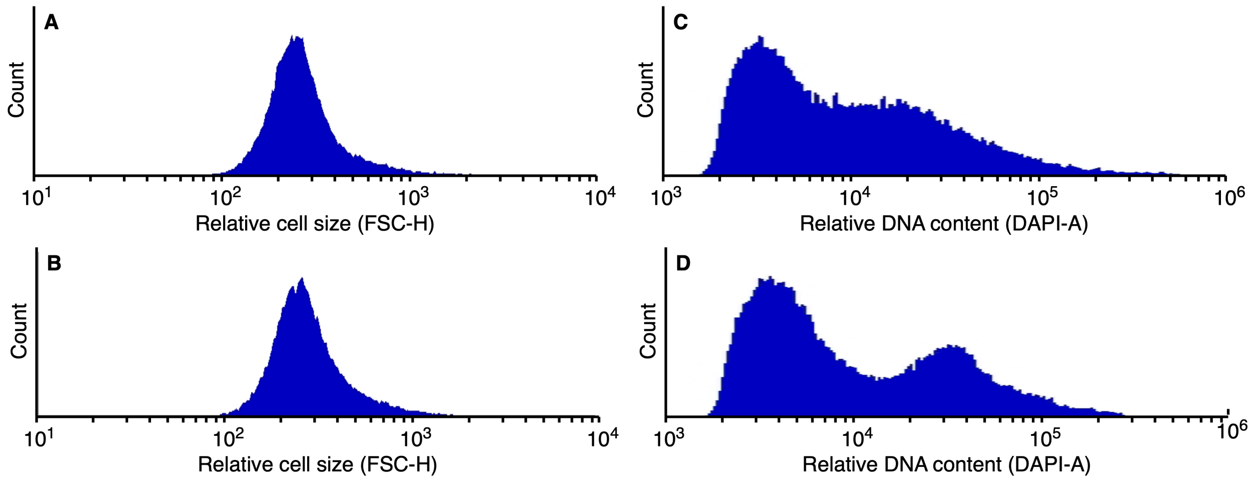


**Figure S18. Cell size and DNA content of *E. meliloti* zone II bacteroids.** (**A**-**B**) Histograms summarizing the distribution of flow cytometry side scattering values of heat-killed *E. meliloti* Rm2011 (**A**) and *E. meliloti* FSM-MA (**B**) zone II bacteroid populations. (**C**-**D**) Histograms summarizing the distribution of flow cytometry DAPI fluorescence values of heat-killed *E. meliloti* Rm2011 (**C**) and *E. meliloti* FSM-MA (**D**) zone II bacteroid populations. Graphs are based on 50,000 cells.

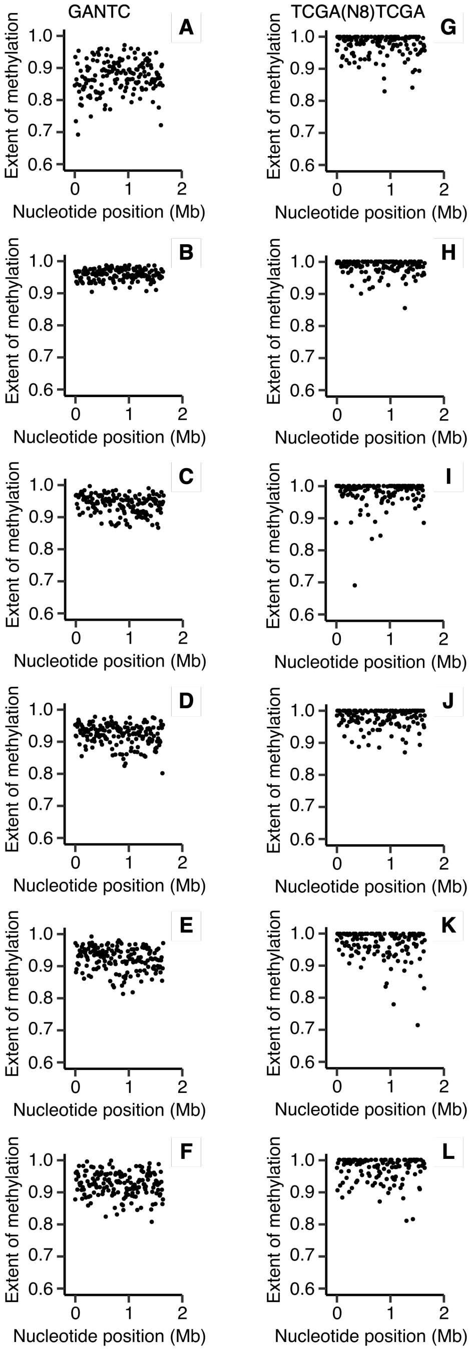


**Figure S19. pSymB-wide DNA methylation of *E. meliloti* FSM-MA bacteroids purified from *M. truncatula dnf* mutant nodules.** The extent of methylation of (**A**-**F**) GANTC and (**G**-**L**) TCGA($N_8$)TCGA motifs across the *E. meliloti* FSM-MA pSymB replicon is shown using a 10 kb sliding window. (**A**,**G**) Bacteroids isolated from *M. truncatula dnf1* mutant nodules. (**B**,**H**) Bacteroids isolated from *M. truncatula dnf5* mutant nodules. (**C**,**I**) Bacteroids isolated from *M. truncatula dnf2* mutant nodules. (**D**,**J**) Bacteroids isolated from *M. truncatula dnf7* mutant nodules. (**E**,**K**) Bacteroids isolated from *M. truncatula dnf4* mutant nodules. (**F**,**L**) Bacteroids isolated from wild-type *M. truncatula* A17 nodules.

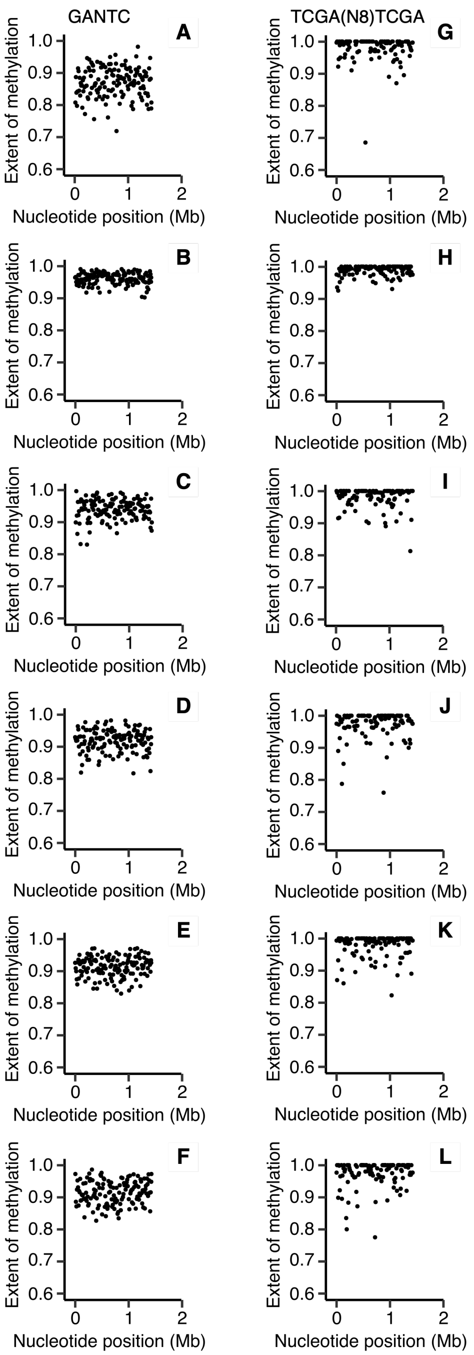


**Figure S20. pSymA-wide DNA methylation of *E. meliloti* FSM-MA bacteroids purified from *M. truncatula dnf* mutant nodules.** The extent of methylation of (**A**-**F**) GANTC and (**G**-**L**) TCGA($N_8$)TCGA motifs across the *E. meliloti* FSM-MA pSymA replicon is shown using a 10 kb sliding window. (**A**,**G**) Bacteroids isolated from *M. truncatula dnf1* mutant nodules. (**B**,**H**) Bacteroids isolated from *M. truncatula dnf5* mutant nodules. (**C**,**I**) Bacteroids isolated from *M. truncatula dnf2* mutant nodules. (**D**,**J**) Bacteroids isolated from *M. truncatula dnf7* mutant nodules. (**E**,**K**) Bacteroids isolated from *M. truncatula dnf4* mutant nodules. (**F**,**L**) Bacteroids isolated from wild-type *M. truncatula* A17 nodules.

**Dataset S1 (separate file).** Cell cycle regulated genes belonging to transcripts with at least one GANTC site situated within the 125 bp upstream region. The first column indicates the gene locus tag, while the second column indicates the cell cycle expression group of the gene as defined by De Nisco et al (2014) (6).
